# Reliability of web-based affective auditory stimulus presentation

**DOI:** 10.1101/2021.02.08.430267

**Authors:** Tricia X.F. Seow, Tobias U. Hauser

## Abstract

Web-based experimental testing has seen an exponential growth in psychology and cognitive neuroscience. However, paradigms involving affective auditory stimuli have yet to adapt to the online approach due to concerns about the lack of experimental control and other technical challenges. In this study, we assessed if sounds commonly used to evoke affective responses in-lab can be used online. Using recent developments to increase sound presentation quality, we selected 15 commonly used sound stimuli and assessed their impact on valence and arousal states in a web-based experiment. Our results reveal good test-retest reliability and good internal consistency, with results comparable to in-lab studies. Additionally, we compared a variety of previously used unpleasant stimuli, allowing us to identify the most aversive of these sounds. Our findings demonstrate that affective sounds can be reliably delivered through web-based platforms, which helps facilitate the development of new auditory paradigms for affective online experiments.

## Introduction

Cognitive psychology and neuroscience researchers are increasingly turning towards online worker platforms such as Prolific (https://prolific.ac/) and Amazon’s Mechanical Turk (MTurk; https://www.mturk.com/) for recruiting participants to complete research studies (Stewart et al., 2017). Given the recent COVID-19 pandemic (Wang et al., 2020), online experimental studies may now even be necessary to circumvent restrictions in carrying out in-lab testing. Web-based testing is attractive not only for its convenience, but it also offers a wealth of advantages ranging from its cheap and rapid data collection system to reach large sample sizes necessary for well-powered research (Gillan & Daw, 2016) to the availability of more diverse or under-represented populations (Berinsky et al., 2012; Casey et al., 2018; Goodman et al., 2013; Levay et al., 2016; Majima et al., 2017; Shapiro et al., 2013). Moreover, there is strong evidence that web collected data is qualitatively on par with data collected from traditional participant pools (Berinsky et al., 2012; Klein et al., 2014; Paolacci et al., 2010), with high internal reliability and test-retest reliability (Shapiro et al., 2013). Pioneering webbased studies across cognitive neuroscience, political science and mental health have indeed produced impactful and replicable findings (Rollwage et al., 2018; Rouault et al., 2018; Schulz et al., 2020; Seow & Gillan, 2020).

However, not all fields have enthusiastically adopted the online approach owing to several criticisms of web-based testing, such as the lack of experimental control. Experiments which utilize auditory stimuli must contend with limited control over audio volume adjustment as system sound settings are not accessible through the standard internet browser as a necessary security measure. Additionally, it is difficult to certify a consistent quality of sound presentation across participants due to the variability in participants’ audio delivery equipment and distractions in the listening environment. While the former issue is difficult to overcome, ensuring that participants wear headphones (including in-ear varieties, i.e., earphones) can help to increase sound quality and reduce interfering noises from the surrounding environment. Recently, headphone screening tests have been developed and validated online (Milne et al., 2020; Woods et al., 2017), allowing an increase web-based audio presentation quality.

Building on this important work, we were interested in testing the viability of sounds presented online to evoke affective responses. Affective stimuli like loud, unpleasant noises are commonly used in cognitive paradigms, for example to induce aversive states and reactions (Neumann & Waters, 2006; Oyarzún et al., 2012; Zald & Pardo, 2002). These paradigms are widely utilized as behavior in these tasks are well-characterized with neural correlates (Büchel & Dolan, 2000; Zald & Pardo, 2002) and computational modelling (Malaka, 1999; Moutoussis et al., 2008; Tzovara et al., 2018), and are even central in investigating psychiatric disorders (Birbaumer et al., 2005; Duits et al., 2015; T. U. Hauser et al., 2016). Yet, these tasks have not been adapted online (except one which uses an alternative method manipulating contexts to induce negative states (Wise & Dolan, 2020)) as it is unknown if affective sounds delivered through the web browser would be able to reliably and effectively evoke the expected emotional responses.

The second intention of the study was to examine which aversive sound stimulus would be most suitable for inducing negative affective responses. Known unpleasant noises vary greatly in semantic category, ranging from female screams (Lau et al., 2008; Morriss et al., 2016), metal screeches (Neumann & Waters, 2006; Zald & Pardo, 2002), high frequency tones (Mirz et al., 2000; Zald & Pardo, 2002), or a loud blast of white noise (Bacigalupo & Luck, 2018; Morris et al., 2001). Unpleasant noises chosen for use in previous research were often either selected by researchers a *priori*, picked from unpublished pilot studies, or chosen from sound databases with affective ratings collected from in-lab sessions that can afford a decent degree of experimental control (Bradley & Lang, 2007; Yang et al., 2018). As such, it is unknown how the valences of these previously utilized aversive stimuli compare against one another and if biases in valence would be similar when presented online with less stringent experimental control.

Here, we capitalize on recent advances in auditory research and sound delivery measures using a headphone screening test and other technical adjustments to optimize online presentation of auditory stimuli. Our aims were two-fold: to (i) investigate if affective sounds presented through an online platform can garner reliable affective responses and (ii) to identify an effective aversive stimulus by comparing the valence of previously utilized aversive noises. We conducted the study with a web-based task where N = 84 participants rated 15 different sounds repeatedly along valence and arousal dimensions. Participants also completed psychiatric questionnaires on anxiety and obsessive-compulsive symptoms. Overall, we found that the affective ratings had good reliability in several measures, good internal consistency, were comparable to ratings from their original sound databases and were not associated with psychiatric severity scores. The most unpleasant sound in our array was a modified female scream (Morriss et al., 2015, 2016, 2020). Our findings suggest that despite limitations of audio experimental control in web-based testing, affective responses can be reliably evoked in participants, validating the use of affective sounds for online research.

## Materials and Methods

### Participants

Data were collected online using Prolific (https://www.prolific.co/) (N = 100). Recruited participants were aged between 18 to 40 years old (mean (*M*) = 26.85, standard deviation *(SD)* = 5.96). We decided to not include older participants because declining sensitivity to high frequency tones spreads to low frequency tones from 40 years onwards (Moore et al., 2014). All individuals were residents of UK and reported normal hearing. The latter meant that they had no past or current personal history of auditory/hearing difficulties including tinnitus, hearing sensitivity (e.g., hyperacusis), hearing loss, use of hearing aid, and current ear infections/inflammation. All participants provided informed consent online after reading the study information and consent pages. Participants were given 30 mins for task completion and were reimbursed at £8.25/hour. All experimental protocols were conducted in accordance with guidelines approved by the UCL Research Ethics Committee (project ID number 15301\001).

### Exclusion criteria

Several pre-defined exclusion criteria were applied to ensure data quality. Participants were excluded if i) their final chosen upper frequency threshold (see *<u>Sound frequency calibration</u>*) was below 8,000Hz (N = 11) which indicated either faults in the auditory setup or a failure to understand instructions, or if ii) they failed the attention check question (i.e., “Demonstrate your attention by selecting ‘A lot’.”) in the questionnaire section (N = 2). Participants with incomplete datasets due to remote data collection issues were also excluded (N = 3). In total, 16 participants (16%) were excluded, leaving N = 84 participants for analysis. Of the remaining sample, 43 (51.19%) identified as female, 40 (47.62%) as male and 1 (1.19%) as other gender.

### Sound array

The sound bite array of 15 sounds was selected to contain a range of (predominantly) unpleasant and pleasant noises. It was assembled from a variety of sources:

i. 4 sounds from the open-source sound database Expanded Version of the International Affective Digitized Sounds (IADS-E) (Yang et al., 2018): female scream (ID: 0276), cicada (ID: 0335), sea wave (ID: 0921) and a piano melody (ID: 1360),
ii. 4 sounds generated using the sound editor software *Audacity* (http://audacityteam.org/) version 2.4.2: pink noise, Brownian noise, 800Hz sine tone and 5,000Hz sine tone, all with 1500ms duration at 0.8 amplitude,
iii. 1 sound from a collaborative database of Creative Commons Licensed sounds, *Freesound* (https://freesound.org): a dentist drill (https://freesound.org/people/alexanderwendt/sounds/385680/),
iv. 3 sounds from previous studies of aversive learning: white noise (Bacigalupo & Luck, 2018), a modified female scream (Morriss et al., 2015, 2016, 2019) (which was altered from the original sound in the second version of the International Affective Digitized Sounds database (IADS-2) (Bradley & Lang, 2007), ID: 277), high frequency tone and a metal screech (Zald & Pardo, 2002), and
v. 2 sounds of 2000ms high frequency tones generated in-browser from frequencies based on participants’ input (see *<u>Sound frequency calibration</u>*) with *react-tone* (https://www.npmjs.com/package/react-tone) version 1.1.1.

All sounds were cut to 2000ms or 1500ms in *Audacity* if the original file was longer.

### Procedure

The experimental task was programmed with *ReactJS* and bootstrapped with *Create React App* (https://create-react-app.dev/). The use of headphones (includes earphones) was required in this study. After participants provided online consent, participants were directed to adjust their volume settings, which was intended to help avoid presentation levels that would result in uncomfortably loud or inaudible sounds, before performing headphone check test. Once through, participants were told to indicate their maximum audible frequency level that would subsequently inform two sounds of the sound array (see **<u>Sound array</u>**). Participants then rated all sounds along valence and arousal dimensions, and completed three psychiatric questionnaires. Details of the procedures are further detailed below:

#### Volume calibration

Volume intensity depends on both the computer system and browser sound settings. As the former cannot be altered by the experimenter, we relied on adjustments of the browser-based volume to manipulate loudness. To do this, we first instructed participants to adjust their computer sound setting to 30% of the system maximum. Thereafter, participants were presented with a noise sample which could be adjusted in volume via browser sound settings from 0 to 100 along a logarithmic scale of lower bound 1 and higher bound 100 (as changes of perceived hearing is better described along a logarithmic scale). Participants were told to adjust the indicator of the scale (initial value = 80) until the sound bite volume was loud but comfortable, allowing as many repeated sound presentations as needed. The final set volume level was subsequently used for the rest of the task.

#### Headphone screening test

We required participants to use headphones as this improves the control of sound delivery and attenuates environmental interference. Participants needed to pass a headphone check task (Woods et al., 2017) to ensure that they were wearing headphones. The task consisted of 6 questions where they had to discriminate the sound intensity of several tones (i.e., which was the quietest) where one was presented with a phase difference of 180° between stereo channels. The sound volumes are difficult to discriminate with loudspeakers (as phase-cancellation is imperfect), but easy with headphones. Sound stimuli of the headphone check task were presented in a randomized order. Participants were required to answer at least 5/6 of the questions correctly to proceed to the next stage, otherwise they were directed back to the volume adjustment module to calibrate their sound settings before attempting the headphone test again. Most participants passed the headphone check on the first attempt (N = 68, 80.95%), while 15.48% of the participants took the test twice (N = 13), and the reminder (N = 3, 3.57%) took more tries, up to a maximum of five attempts.

#### Sound frequency calibration

This part of the experiment was intended to determine the individual’s upper frequency threshold, which was subsequently used to select high frequencies as unpleasant noise stimuli. Sensitivity to high frequencies range tends to decline with age (Lee et al., 2012; Moore et al., 2014), and as such we sought to determine the maximum audible frequency for each individual which we could then derive subjectively high but audible frequency tones as part of the sound array. In this section, participants were shown a logarithmic (as frequency perception follows a logarithmic perception) scale that represented frequency of a sine tone, with the indicator at an initial frequency of 8,000Hz. They were told to adjust the frequency of the tone three times (first scale: ranged 5,000Hz to 20,000Hz, second and third scales: 1,000Hz less from previous chosen frequency to 20,000Hz; all logarithmic) to a pitch where they could just hear the tone. We did not find a significant influence of age on the final chosen frequency level in our relatively narrow participant age range *(**Supplementary Fig S1**).* Two frequencies were derived from this upper frequency threshold which were subsequently used as part of the sound bite array for the ratings task—one 3/4 (Frequency 1) and one ½ (Frequency 2) of the chosen frequency level set on a logarithmic 10,000Hz to 20,000Hz scale.

#### Auditory ratings

The participants rated each of the sounds on two affective scales: valence and arousal. These dimensions are considered classic, primary dimensions of emotion that account for most of the variance in emotional judgments (Bradley & Lang, 1994) and were measured in prior rating studies (Bradley & Lang, 2007; Yang et al., 2018). We used 0 to 100 continuous scales. For valence, the scale ranged from very unpleasant (0) to neutral (50) to very pleasant (100), while the arousal scale ranged from very sleepy (0) to neutral (50) to very awake (100). Each of the 15 sound bites were presented 4 times (60 times in total); twice at the same volume set at the beginning (100%) (see *<u>Volume calibration</u>*), and twice more at half the logarithmic scale (lower bound 1 and higher bound 100) of the original volume setting (50%). This was intended to enable test-retest reliability analyses of comparing affective ratings across the repeated presentations of the specific sound stimuli. Sounds were presented in a randomized order.

The default value indicator for both affective scales were also randomized to begin between 35 and 65 for every presentation. Participants were required to adjust the slider on both scales before they could move on to rate the next sound. The participants had the option to play the sounds multiple times during rating, but sounds were generally played once (*M* per participant = 1.09, *SD* = 0.16). Participants could also indicate with a checkbox if they could not hear the sound; none of the sound presentations for any participant were signaled as such.

#### Questionnaires

Lastly, participants provided basic demographic data (age and gender) and completed three self-report psychiatric questionnaires. We administered questionnaires assessing symptoms of state (STAI-Y1) and trait anxiety (STAI-Y2) using both scales of the State-Trait Anxiety Inventory (Spielberger et al., 1983) as well as obsessive-compulsive disorder (OCD) symptoms using the Obsessive-Compulsive Inventory - Revised (OCI-R) (Foa et al., 2002). The presentation order of the questionnaires was randomized.

### Analyses

All analyses were conducted in R, version 3.6.0 via RStudio version 1.2.1335 (http://cran.us.r-project.org). All mixed effect models were estimated with the *lmer()* function from the *lme4* package with the *lmerTest* package for statistical tests. Linear regression models were performed with *lm()* function from the *stats* package. For correlation tests, we used non-parametric Spearman’s correlation tests with no ties to account for the non-normality of the data, which were conducted with the *cor.test()* function from the *stats* package. The internal consistency measure, Cronbach’s alpha, was estimated with the *alpha()* function from the *psych* package. Paired t-tests were calculated with the *t.test()* function from the *stats* package.

### Reliability measures

1. <u>*Intra-participant test-retest reliability.*</u> We tested whether participants’ ratings were reliable by correlating the ratings (valence or arousal) of the first presentation of each individual sound with the rating of its identical, repeated presentation (same volume level) for each participant. We then tested if volume *(Volume:* low (50%) or high (100%)) or affective scale type *(ScaleType:* valence or arousal) were linked to this reliability correlation measure *(ParticipantReliability).* Both *Volume* and *ScaleType* were taken as factors. We used a mixed-effects model in which *Volume, ScaleType* were fixed effect covariates, with them and the intercept taken as random effect predictors. The model was: *ParticipantReliability* ~ *Volume + ScaleType* + (1 + *Volume + ScaleType | Subject).*
2. <u>*Inter-participant test-retest reliability.*</u> We also asked if there were inter-participant differences in their ratings across the repeated sound presentations. For this, we calculated the means and standard deviations of all affective ratings by volume and affective scale type for each participant for the sounds’ first and second presentation separately. We then examined the correlation between the means and standard deviations obtained across the two presentations.
3. <u>*Intra-sound test-retest reliability.*</u> The sounds themselves could also differ in test-retest reliability variance. To investigate this, we correlated the affective ratings from the two repeated presentations across all participants for each sound at its unique volume level. We additionally tested if *Volume* and *ScaleType* influenced the sound reliability *(SoundReliability)* with a mixed-effects model replacing *ParticipantReliability* in the prior analysis.
4. <u>*Internal consistency.*</u> Internal consistency of the affective ratings were measured in prior studies (Bradley & Lang, 2007; Yang et al., 2018), which reflected the similarity of the ratings across the sounds. Likewise, we measured the consistency of the ratings using Cronbach’s alpha for all sound presentations for both valence and arousal scales.
5. <u>*Correlation with prior ratings.*</u> To examine the robustness of our ratings which were garnered online versus in-lab rating studies, we compared the affective ratings of 5 sounds from the current online study (the averaged rating across repeated presentation at 100% volume) with the ratings from their original lab-based study (Bradley & Lang, 2007; Yang et al., 2018). We used Pearson’s correlation in this measure as we were interested in the numerical rather than ordinal relationship between the ratings.

### Ranking affective ratings

We ranked the sounds in order of valence and arousal ratings. For these analyses, we used the averaged rating across repeated presentations of the same sound at its specific volume. To test for significance in rating differences between sounds, we conducted paired t-tests. We also asked if affective ratings were influenced by the volume of the presented sound by testing if the affective ratings *(ScaleRating)* were associated with *Volume* with a mixed-effects model for both affective scale types. The model was: *ScaleRating* ~ *Volume* + (1 + *Volume | Subject).*

### Unique high frequency tones

High frequency tones are known to be unpleasant (Zald & Pardo, 2002). However, upper hearing frequency limits might differ for participants given the effect of age on hearing thresholds (Lee et al., 2012). Though we attempted to circumvent this issue with our age exclusion criteria (≤40 years), we also allowed participants to identify subjectively high frequencies for themselves (Frequency 1 and Frequency 2; see *<u>Sound frequency calibration</u>*) in addition to having two separate sine wave stimuli with frequencies set at 800Hz and 5,000Hz. As such, unique frequencies were played across the participants for Frequency 1 and Frequency 2 in the sound array. We thus tested if the unique frequencies that participants heard *(Frequency*, z-scored) influenced their affective ratings *(ScaleRating)* for valence and arousal dimensions with the model: *ScaleRating* ~ *Frequency* + (1 + *Frequency | Subject).*

### Questionnaires

Finally, we asked if the affective ratings *(ScaleRating)* or intraparticipant rating reliability *(ParticipantReliability)* were associated with the psychiatric symptom severity *(QuestionnaireScore*, z-scored) and if that main effect was modulated by *Volume* for both valence and arousal dimensions. For this, we used a mixed effects model: *ScaleRating* ~ *QuestionnaireScore* * *Volume* + (1 + *Volume | Subject)* and a linear model: *ParticipantReliability* ~ *QuestionnaireScore* * *Volume.* Also see ***Supplementary Fig. S4*** and ***Supplementary Fig. S5*** for questionnaire score distributions and correlations respectively.

## Results

### Volume and frequency adjustment

#### Volume adjustment

First, to ensure that sounds presented in this study were appropriate (i.e., not dangerously loud), we let participants calibrate the browser sound volume to a level that was comfortable for them (see *<u>Volume calibration</u>).* We subsequently used the adjusted sound volume from the successful attempt of the headphone task (see *<u>Headphone screening test</u>*) for the rest of the experiment. Participants generally chose a volume level on the scale that was close to the default of 80 (mean *(M)* = 76.49, standard deviation *(SD)* = 16.97), with a range of 27 to 100 *(**Fig. 1a**)*.

**Fig. 1.**
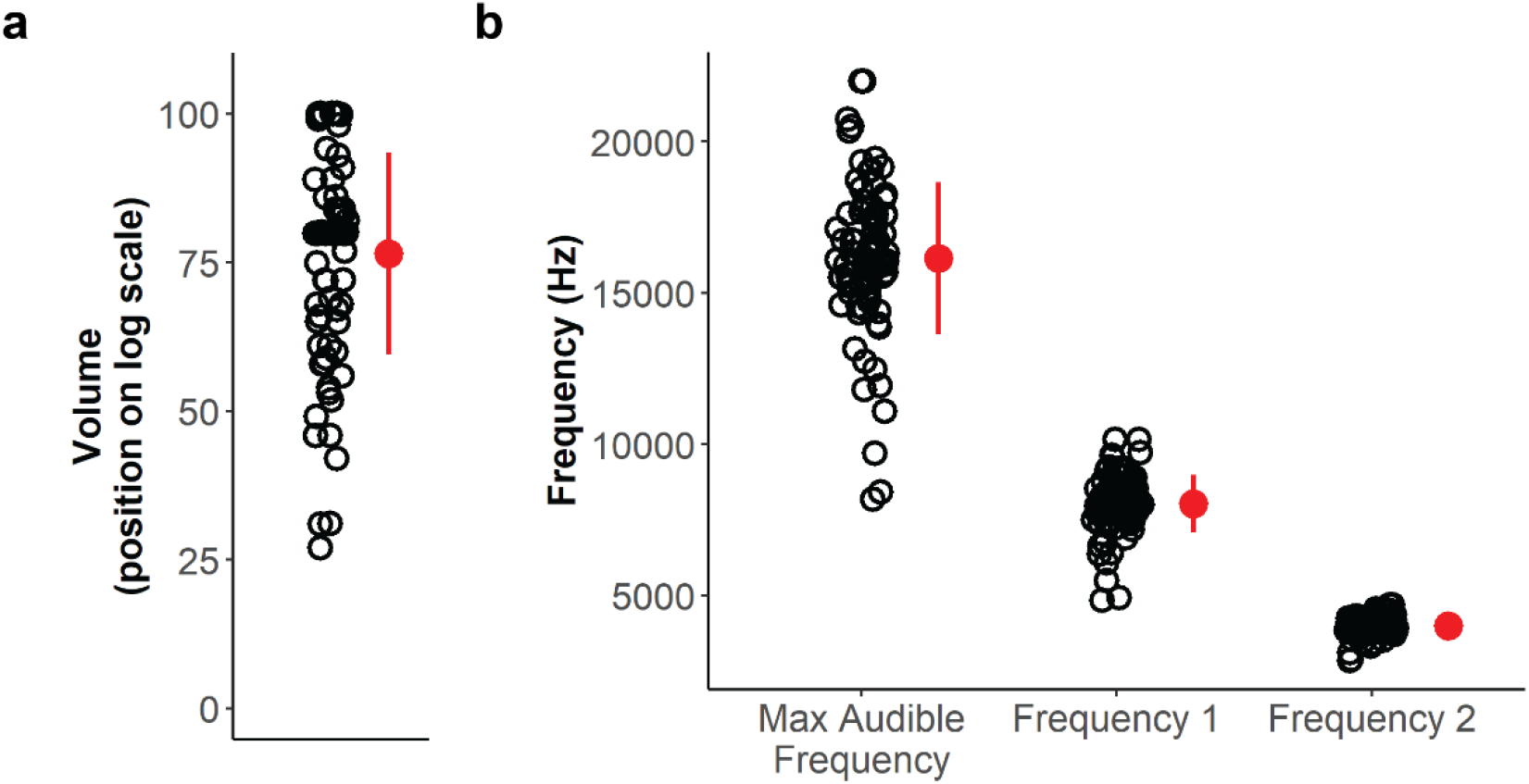
Adjusted volume and frequency levels. ***a*** Distribution of adjusted volume of browser sound settings. Participants were instructed to keep their computer system sound settings at 30% of the maximum before calibrating the browser volume on a (log) scale to a level that was appropriate (not uncomfortably loud) for them. ***b*** Distribution of the maximum audible frequency selected, and the frequencies derived to use in the sound array for subsequent rating which were unique to each participant. Frequency 1 was derived from 70% (log-scaled), and Frequency 2 was 50% (log-scaled), of their threshold frequency. Circles in the graphs represent volume/frequency level per participant. Red marker indicates mean and error bars indicate standard deviation.

#### Frequency adjustment

Next, we attempted to identify frequency thresholds for each participant in order to select subjective high frequency pitches for each participant as an unpleasant noise. Participants chose 16,141.15Hz on average (*SD* = 2,516.79Hz, ranging between 8,184Hz to 22,000Hz) *(**Fig. 1b**)* as the pitch where they could just hear the tone. These frequency levels are line with hearing threshold studies where the 22-35 age group sample is thought to have approximately 16,000 Hz (at 63.03 dB, a relatively moderate to loud volume) as their upper frequency threshold (Lee et al., 2012). For the ratings task, we used sine tones of frequency levels which were slightly lower than their self-reported threshold for each participant (see *<u>Sound frequency calibration</u>)—*Frequency 1 which was 70% (log-scaled) of the threshold (*M* = 8,033.45Hz, *SD* = 959.89Hz, min = 4,838.65Hz, max = 10,158.21Hz), and Frequency 2 which was 50% (log scaled) of the threshold (*M* = 4,004.45Hz, *SD* = 326.87Hz, min = 2,860.77, max = 4,690.42Hz).

### High reliability of affective ratings

#### Intra-participant test-retest reliability

In this study, we sought to examine the reliability of the affective ratings by our online participant pool. To do so, each of the sounds (at two volume levels; 50% and 100%) was played twice to examine how similar the same sound was rated across two presentations. For this, we calculated the Spearman correlation *(ρ)* between the two ratings of the same sound played for each participant at its specified volume level. We found that the correlations across the ratings of these repeated sound presentations were positive and high for across all participants in both valence (*M* = 0.89, *SD* = 0.08) and arousal (*M* = 0.79, *SD* = 0.14) *(**Fig. 2a**)*, showing good test-retest reliability. We generally observed that valence ratings had a stronger positive correlation between repeated presentations than arousal ratings *(β* = 0.11, *SE* = 0.02, *p* < 0.001) *(**Fig. 2a**)*, indicating that participants gave more reliable valence than arousal ratings. There was no significant impact of volume on correlation strength between the repeated ratings *(β* = 0.02, *SE* = 0.01, *p* = 0.10). Overall, these findings suggest that participants rated sounds very similarly across their repeated presentation on both valence and arousal scales, presenting the ratings as a reliable measure of emotional response.

**Fig. 2.**
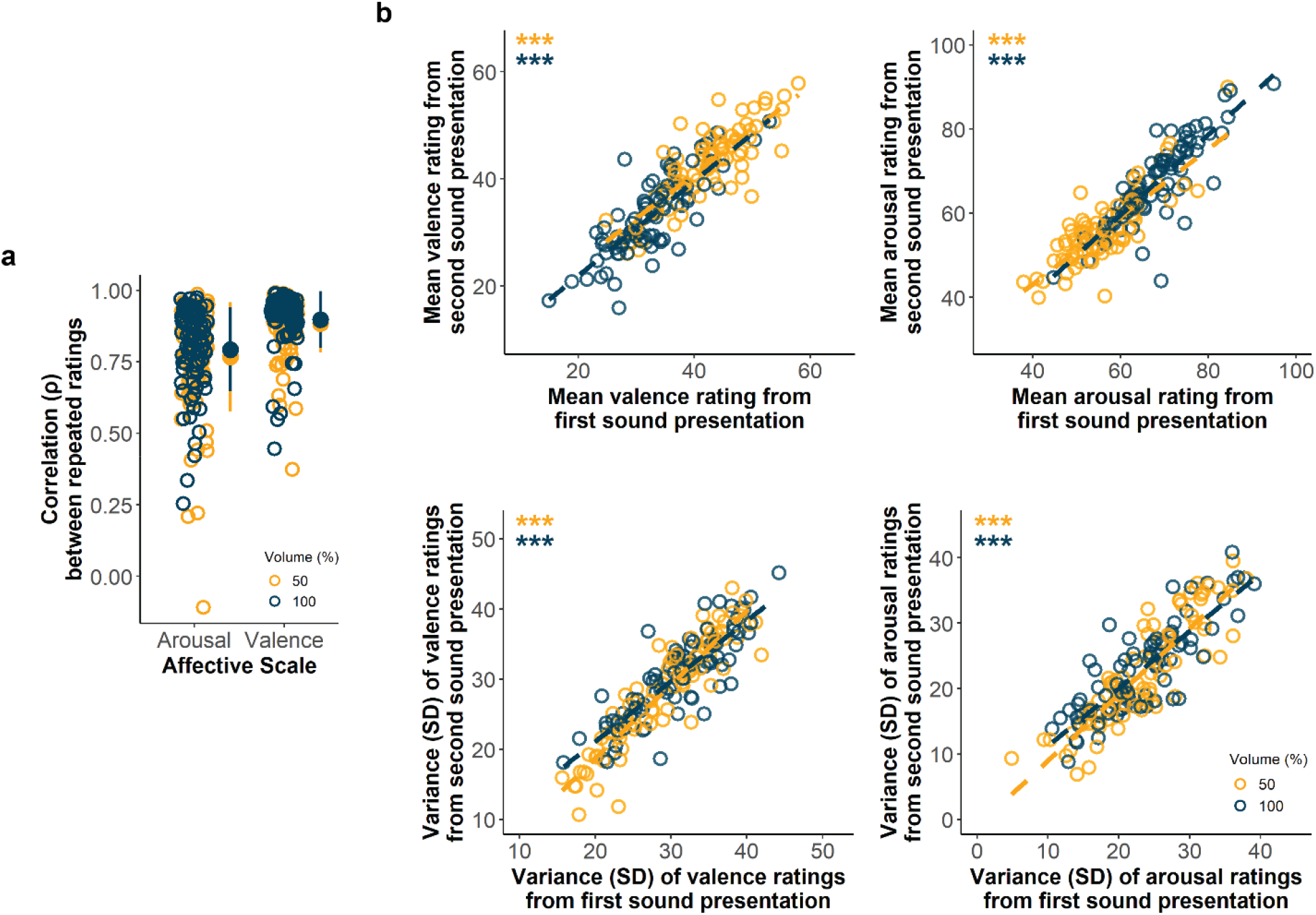
Intra- and inter-participant reliability of affective ratings. ***a*** Distribution of intra-participant reliability. Affective ratings for each sound were correlated with those from its repeated presentation as the intra-participant reliability measure. These were found to be high for both arousal and valence in both volumes. Circles represent Spearman’s rank correlation estimate ρ across repeated sound ratings for each participant for a specified volume/scale. Marker indicates mean and error bars indicate standard deviation. ***b*** Inter-participant reliability as measured by the comparison of the means and standard deviations of all sound ratings in the first (x-axis) versus second (y-axis) stimulus presentation. Strong positive correlations between the measures for both arousal and valence indicate reliable rating dynamics. Circles represent the mean/standard deviation for each participant at its specified volume, with the dashed line indicating the linear relationship between means/standard deviations. ***p < 0.001 (Spearman’s correlation).

#### Inter-participant test-retest reliability

Next, we investigated how similarly participants provided their ratings in terms of their overall rating means and standard deviations across the two sound presentations. We found that participants gave comparable rating dynamics over the repeated presentations in both valence and arousal scales— correlations between the mean rating of all sounds between the first and second sound presentation across all participants were high (valence: *ρ =* 0.87, *p* < 0.001; arousal: *ρ =* 0.87, *p* < 0.001) and the variances were also very similar (valence: *ρ =* 0.88, *p* < 0.001; arousal: *ρ =* 0.86*, p* < 0.001) *(**Fig. 2b**).* Again, these results support the high reliability of participants providing affective ratings for the sounds presented online.

#### Sound-specific test-retest reliability

Across our array of 15 sounds, we also wondered if the sounds intrinsically differed in reliability of their ratings. As such, we examined the degree of correlation between the repeated ratings across all participants for each sound separately. Overall, all sounds were able to reliably induce affective ratings at both 50% (*M* = 0.68, *SD* = 0.06) and 100% volume (*M* = 0.70, *SD* = 0.07) *(**Supplementary Fig. S2**).* At 100% volume, the sound which gave the most reliable ratings was the modified female scream of Morriss *et al. (ρ =* 0.78, *p* < 0.001) while the least reliable was the Pink Noise stimulus *(ρ =* 0.56, *p* < 0.001) *(**Supplementary Fig. S2**)*.

#### Internal consistency

Prior studies have also reported good internal consistency of sound ratings both valence and arousal affective dimensions (Yang et al., 2018). Similarly, we estimated Cronbach’s alpha for all sounds per scale type. The top seven valence sounds (100% and 50% volume for Cicada, Sea Wave and Piano Melody, and 50% volume Brownian Noise), which was the same as the bottom seven arousal sounds, were automatically identified as being negatively corelated with its respective scale type. This indicated that our sound array had a balanced range of sounds with differing valence and arousal. The internal consistencies were found to be high for both scale types after automatic sign reversion of those sounds in the estimation (valence: α = 0.93, arousal: α = 0.94).

#### Correlation with prior ratings

Finally, we asked if the ratings we collected online were comparable to those from in-lab studies to test if the decreased degree of experimental control of web-based audio delivery impacted the affective ratings. We gathered the ratings of 5 sounds in our stimuli array (played at 100% volume) to compare them with those from their original studies which were all conducted in traditional laboratory environments (Bradley & Lang, 2007; Yang et al., 2018) Pearson’s correlation of the sounds across ours and the prior studies for both affective dimensions were high (arousal: *r* = 0.98, *p* = 0.003; valence *r* = 0.96, *p* < 0.01) *(**Fig. 3**).* These results suggest that the affective ratings we collected online were reliable and valid, and rated similarly to those measured in well-controlled in-lab environments.

**Fig. 3.**
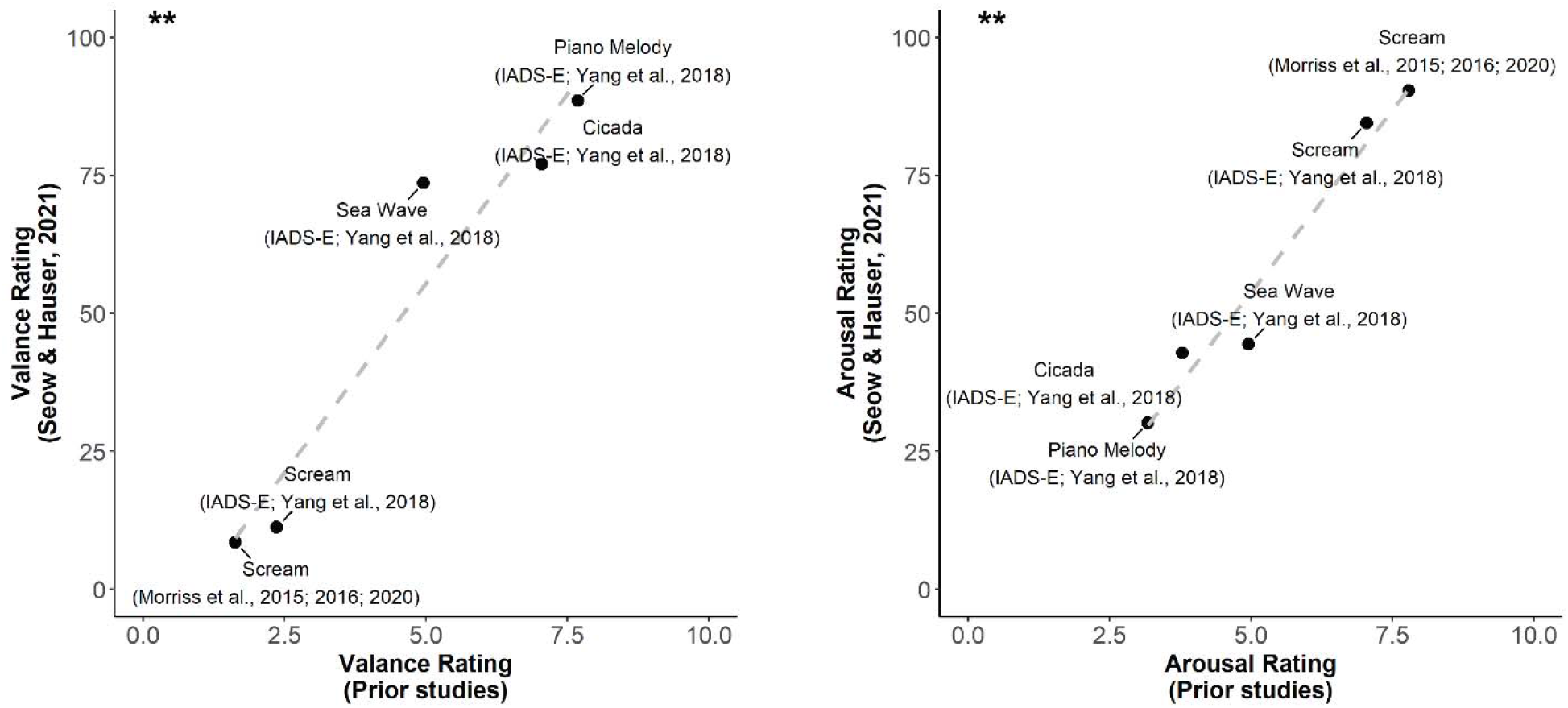
Rating comparison with prior studies. Relationship between current affective ratings and those from prior studies (Bradley & Lang, 2007; Yang et al., 2018). Pearson’s correlation across the 5 sounds in both rating dimensions were high (arousal: r = 0.98, p = 0.003; valence: r = 0.96, p < 0.01), indicating that our ratings collected online are very similar to those from traditional in-lab studies. (n.b. the Morriss et al. scream is a modified version of the original sound bite from the IADS-2 (Bradley & Lang, 2007), whose ratings were used for comparison here.) **p < 0.01 (Pearson’s correlation).

#### Ranking of affective ratings

As part of the study, we aimed to identify the most unpleasant sound amongst a variety that have been utilized in prior research. Female screams were found to be the most unpleasant, with the modified female scream of Morriss *et al. (M* = 8.48, *SD* = 12.09) being rated as significantly more unpleasant than the scream from the IADS-E database (Yang et al., 2018) (*M* = 11.24, *SD* = 12.68) (at 100% volume for both sounds: *t* = −1.99, 95% CI = [−5.52, −0.008], *p* < 0.05) *(**Fig. 4**).* This was followed by high frequency tones, metal sounds, complex noises, and finally natural pleasant noises. The most pleasant sound was the Piano Melody (*M* = 88.57, *SD* = 10.88), which was rated as significantly more pleasant than Cicada sounds (*M* = 81.57, *SD* = 13.81) (at 100% volume for both sounds: *t* = 4.01,95% CI = [4.39, 13.02], *p* < 0.001) *(**Fig. 4***).

**Fig. 4.**
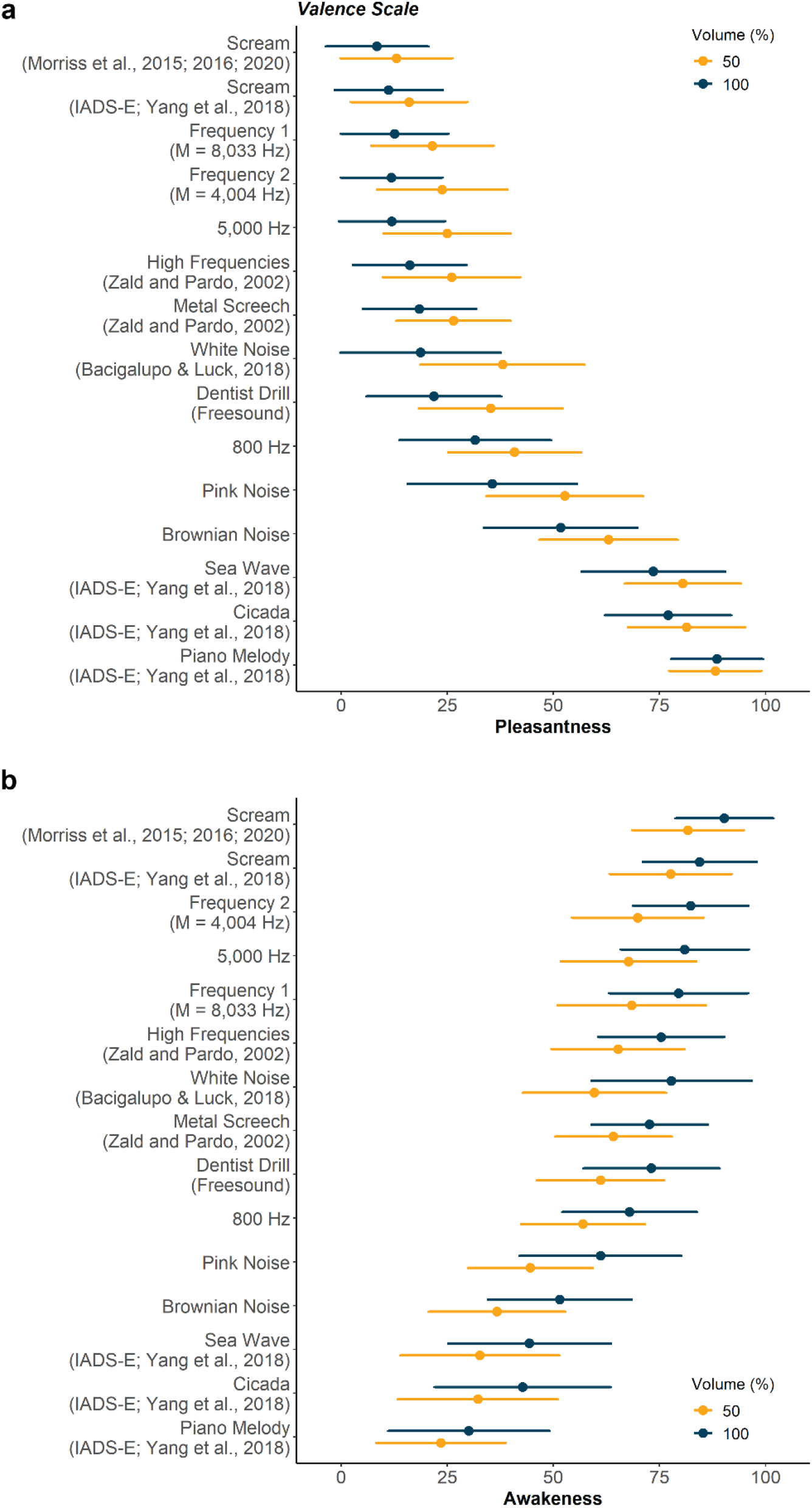
Mean affective ratings for the sound array across all participants. We ranked the affective ratings from the **a** valence and **b** arousal scales to identify the most unpleasant sound in our array, split by volume. The Morriss et al. scream was found to be the most unpleasant and arousing sound. Frequency 1 and Frequency 2 were unique for every individual, based on their calibrated frequency threshold (See <u>Sound frequency calibration</u>), but the mean of the frequencies across all participants for those sounds are indicated in their labels. Marker indicates rating means and error bars indicate standard deviations for all sounds. Sounds are ranked by the averaged rating across volumes. See **Supplementary file 2** for these data.

We also observed that the ratings in valence and arousal dimensions for the current sound array were negatively correlated (Spearman: *p:* = −0.83, *p* < 0.001) *(**Supplementary Fig. S3**).* As such, the sounds followed the almost the same (inverted) ranking pattern for arousal ratings. The Morriss *et al.* modified female scream was identified as the most arousing (*M* = 90.35, *SD* = 11.51), being more arousing than the IADS-E female scream (*M* = 84.50, *SD* = 13.34) (at 100% volume for both sounds: *t* = 4.42, 95% CI = [3.21, 8.48], *p* < 0.001) *(**Fig. 4***). On the other hand, the 50% volume Piano Melody was rated as the least arousing sound (*M* = 23.54, *SD* = 19.03), being significantly lower than the Cicada sound (*M* = 32.23, *SD* = 18.81) (at 50% volume for both sounds: *t* = 3.51,95% CI = [1.75, 6.33], *p* < 0.001).

Overall, the ratings suggest that the Morriss *et al.* modified female scream may be the most ideal aversive stimuli as it was ranked as the most unpleasant and arousing sound amongst others.

#### Volume intensity increases arousal and unpleasantness

High sound intensity is often a factor in inducing aversive responses (Liberman et al., 2006; Neumann et al., 2008), and thus we sought to examine how volume influenced affective ratings. We found that both valence and arousal ratings were modulated by volume. Louder sounds were generally rated as having lower valence (unpleasant) *(β* = −11.98, *SE* = 2.33, *p* < 0.001) and being more arousing (awakeness) *(β* = 22.96, *SE* = 1.82, *p* < 0.001). Separate paired t-tests for ratings of every sound between the two volumes indicated that this was true for all stimuli (valence: all *ts* > 3.72, *p*s < 0.001, arousal: *ts* > 4.17, *p*s < 0.001), except Piano Melody in the valence dimension (*t* = −0.49, 95% Confidence Interval (CI) = [-1.88, 1.14], *p* = 0.62) *(**Fig. 4**).* The sound that reported the largest volume-induced difference in both valence (*t* = 11.73, CI = [16.07, 22.63], *p* < 0.001) and arousal (*t* = −10.11, CI = [−21.69, −14.55], *p* < 0.001) ratings between the two volume levels was White Noise. These results suggest that affective ratings of sound stimuli are generally affected by volume intensity, but sounds may intrinsically differ in this influence.

#### Slight variation in high frequencies not associated with degree of unpleasantness

High frequencies tones are known to be unpleasant, and we observed that the 5,000Hz sine tone was rated significantly more unpleasant than the 800Hz in both volumes (100%: *t* = −9.20, CI = [−22.83, −15.36], *p* < 0.001; 50%: *t* = −9.27, CI = [−19.29, −12.48], *p* < 0.001). As part of the sound array, participants were also presented with two high frequency sine tones that were unique to their (self-reported) maximum audible frequency; Frequency 1 which was lower in pitch than their threshold by ¼, and Frequency 2 which was lower by ½. Valence (*t* = 0.53, 95% CI = [−2.03, 3.50], *ρ* = 0.60) and arousal (*t* = −1.96, 95% CI = [−5.75, 0.04], *p* = 0.05) ratings did not significantly differ between these two sounds at 100% volume (also not at 50% volume: *t*s > −1.45, *p*s > 0.15) *(**Fig. 4**).* We considered that because participants heard objectively different frequencies in these sound bites, affective ratings for Frequency 1 and Frequency 2 might have be influenced by the absolute frequency that was chosen by each participant. However, we found no significant association between the ratings and the objective stimulus frequency (valence: *β* = −0.44, *SE* = 0.70, *p* = 0.53; arousal: *β* = −1.07, *SE* = 0.70, *p* = 0.13). Our findings suggest that while high frequencies are generally considered aversive, variation from about 3,000Hz to 10,000Hz (the sample pool’s minimum of Frequency 2 and maximum of Frequency 1 respectively) did not significantly impact valence ratings.

#### Psychiatric symptom scores are not associated with affective ratings nor rating reliability

We measured psychiatric symptom severities of state anxiety (STAI-Y1), trait anxiety (STAI-Y2) and obsessive-compulsive symptoms (OCI-R) to test whether mental health traits influenced the perception or rating of these stimuli, which could have ramifications for experiments with such populations. Neither valence (STAI-Y1: *β* = −0.96, *SE* = 1.89, *p* = 0.61; STAI-Y2: *β* = −0.21, *SE* = 0.03, *p* < 0.001; OCI-R: *β* = −0.96, *SE* = 1.89, *p* = 0.61) nor arousal (STAI-Y1: *β* = 2.49, *SE* = 1.53, *p* = 0.11; STAI-Y2: *β* = −0.96, *SE* = 1.89, *p* = 0.61; OCI-R: *β* = 2.49, *SE* = 1.53, *p* = 0.11) ratings had a significant relationship with the questionnaire scores (***Fig 5a***). There was also no interaction effect of psychiatric traits with volume in either dimension (arousal: STAI-Y1: *β* = −2.03, *SE* = 1.83, *p* = 0.27, STAI-Y2: *β* = −2.03, *SE* = 1.83, *p* = 0.27, OCI-R: *β* = −2.03, *SE* = 1.83, *p* = 0.27; valence: STAI-Y1: *β* = 1.03, *SE* = 2.34, *p* = 0.66, STAI-Y2: *β* = 1.03, *SE* = 2.34, *p* = 0.66, OCI-R: *β* = 1.03, *SE* = 2.34, *p* = 0.66) on the ratings (***Fig 5a***).

**Fig. 5.**
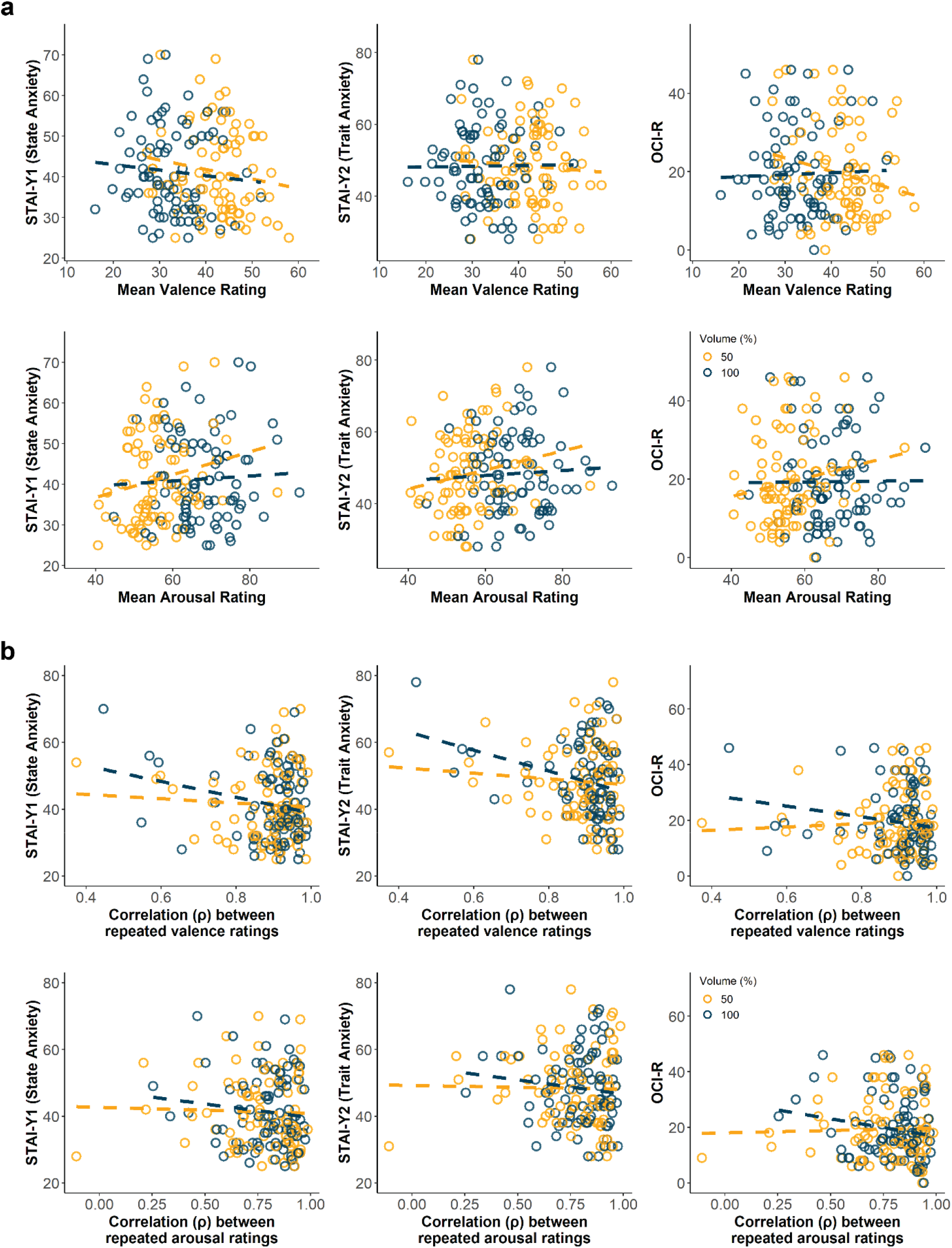
Correlation of questionnaire scores with affective ratings or participant reliability. For illustration purposes, the scatter plots depict the relationship between **a** affective ratings or **b** ratings reliability with questionnaire total scores. None of the correlations were significant, indicating that affective ratings nor its reliability was affected by psychiatric symptom severity. Circles represent either the mean affective rating for all sounds per participant or intra-participant reliability measure, while the dashed lines represent the linear relationship between questionnaire score and mean rating/reliability estimate at its specified volume.

Lastly, we examined if psychiatric traits would be associated to the intra-participant test-retest reliability measure. There was no significant relationship between reliability estimates and any questionnaire scores for arousal (STAI-Y1: *β* = −0.006, *SE* = 0.02, *p* = 0.75, STAI-Y2: *β* = −0.004, *SE* = 0.02, *p* = 0.85, OCI-R: *β* = 0.006, *SE* = 0.02, *p* = 0.76) or valence (STAI-Y1: *β* = −0.008, *SE* = 0.01, *p* = 0.47, STAI-Y2: *β* = 0.006, *SE* = 0.01, *p* = 0.62, OCI-R: *β* = 0.006, *SE* = 0.01, *p* = 0.62) ratings (***Fig 5b***). Similarly, volume did not interact with any of these relationships (arousal: STAI-Y1: *β* = −0.02, *SE* = 0.02, *p* = 0.33, STAI-Y2: *β* = −0.02, *SE* = 0.02, *p* = 0.20, OCI-R: *β* = −0.02, *SE* = 0.02, *p* = 0.15; valence: STAI-Y1: *β* = −0.01, *SE* = 0.03, *p* = 0.71, STAI-Y2: *β* = −0.01, *SE* = 0.03, *p* = 0.63, OCI-R: *β* = −0.03, *SE* = 0.03, *p* = 0.26) (***Fig 5b***). In sum, the psychiatric symptom severities we tested in our participants were not linked to their affective ratings nor reliability of those ratings.

## Discussion

In this study, we examined the feasibility of inducing affective states in webbased online studies using sound stimuli. We find that with the right technical measures in place, we can reliably induce affective states using sound stimuli similar to in-lab studies, which demonstrates that affective audio stimuli can be used well in web-based tasks. Concretely, we found that the ratings were reliable, had good internal consistency and were comparable to those reported in prior studies which collected ratings in controlled in-lab environments. Moreover, we compared several unpleasant sounds consistently and found that a modified female scream led to the most aversive state. Our study thus lays the ground to use affective sound stimuli for inducing negative affective states in online studies, a much needed means for cognitive online studies.

First, we took on the challenge of ensuring good auditory delivery on a webbased platform. In our procedure, participants were first screened to ensure they wore headphones (Woods et al., 2017) to reduce distractions in the listening environment. Thereafter, they were to set their computer system volume at a fixed level, to enable a more consistent sound intensity level across participants, before the final adjustment of the browser volume setting. While we could not objectively track if participants followed our instructions, the high pass rate (80.95%) of the headphone check task with one attempt signaled that participants were generally compliant. Moreover, while there was wide variation in the final browser volume level chosen, majority of the participants selected close to the loud intensity of the default. Lastly, none of the sounds was indicated as inaudible in the rating task, which may occur if too low volumes were picked. Overall, we were confident that our procedure enabled a more consistent and decent presentation of sound quality online.

To examine the test-retest reliability of sound ratings, we specifically designed the task such that participants gave repeated ratings of each sound on two scales, arousal and valence. We found that participants reported very similar ratings across the two presentations of a particular sound, and they also gave comparable rating dynamics in terms of their overall rating means and variances over the repeated presentations. Prior auditory affective rating studies tested in stricter, in-lab environments also examined reliability in terms of internal consistency—likewise, the internal consistency estimates we measured from our ratings were high and similar (e.g. α = 0.95 for valence, α = 0.92 for arousal in (Yang et al., 2018) versus here: α = 0.93 for valence, α = 0.94 for arousal in our study). These results suggest that the affective ratings collected online were highly reliable. More fine-grained analyses show that the sounds intrinsically differed in their ratings reliability, but all of them (at 100% volume) evoked decent reliability of a correlation of *ρ >* 0.60 *(**Supplementary Fig. S2**).* More importantly, as several of our sounds were taken from an open-source sound database (Yang et al., 2018), we were able to compare our ratings that were presented online to ratings that were previously collected in a traditional in-lab setting that were afforded much stricter experimental control. Notably, our sample size was larger than the previous study (N = 84 participants rated each sound versus N = 22 in Yang *et al.)*, affording greater statistical power, and we replicated the ratings of these sounds very closely. Overall, our results support the validity of a web-based presentation for sounds to evoke their expected affective responses.

A second aim of this study was to suss out the most unpleasant sound for use as an effective aversive stimulus. Unpleasant sounds come from a wide range of categories (Bradley & Lang, 2007; Yang et al., 2018), from naturalistic sounds such as female screams to more synthetic sounds like high frequency tones or white noise. Varying sounds have been used in prior research as aversive stimuli to drive learning (Bacigalupo & Luck, 2018; Morriss et al., 2015, 2016, 2019; Zald & Pardo, 2002), but it is unknown how they compare to each other given the different methods of selection. While some sounds have their affective ratings documented in sound databases, others are often chosen after unpublished pilot studies or selected *a priori.* Here, we specifically rated some of these unpleasant sounds together to compare their valence ratings. We found that white noise and metal screeches were mildly aversive, followed by high frequency tones (>4,000Hz), while more naturalistic sounds such as the female scream were rated as the most aversive. The Morris *et al.* scream was rated as the most unpleasant and arousing sound, indicating that it may be the most affective aversive stimulus amongst that we tested.

In this same vein, we manipulated loudness intensity systemically (full and half volume) as the volume is often key in eliciting negative affect (Liberman et al., 2006; Neumann et al., 2008). One previous rating study reported no changes in affective ratings owing to sound intensity differences (Bradley & Lang, 2007), while another noted that volume negatively correlated with valence and positively correlated with arousal (Yang et al., 2018), though this was due to natural volumes in the environment rather than an intended task design. We replicated this same effect with our systematic volume manipulation here. Notably, the sounds intrinsically differed in the strength of influence of volume on their affective ratings—for instance, the ratings for White Noise varied the most depending on volume intensity amongst our array sounds. Nonetheless, volume did not seem to influence valence ratings more so than the type of sound itself *(**Fig. 3**)*, suggesting that these sounds are valanced because of their inherent characteristics, and less because of the intensity they were perceived at.

High frequencies are known to be unpleasant (Mirz et al., 2000; Zald & Pardo, 2002) but upper frequency thresholds are subjective to the individual and are heavily dependent on age (Fozard, 1990; Lee et al., 2012; Moore et al., 2014). We attempted to circumvent this issue by limiting the age in our participant pool to a maximum of 40 years. The final chosen upper frequency threshold levels of our sample did show variation, but this was not related to the participants’ age *(**Supplementary Fig. S1**).* For rating, we presented high frequency tones that were subjective to each participant by utilizing two frequencies that were slightly lower than their unique threshold frequency. In general, we found that the fixed high frequency tone (5,000Hz) was more unpleasant than low frequency (800Hz), but variation in the subjective high frequency tones (from around 3,000Hz to 10,000Hz) did not influence the degree of valence in ratings. Given the lack of relationship between high frequencies and unpleasantness, in addition to some participants having rather low upper frequency threshold levels *(**Supplementary Fig. S1**)*, high frequencies may not be the most decisive aversive stimuli as compared to others like naturalistic screams which were ranked as more unpleasant and are straightforward to use.

Lastly, we collected several psychiatric symptom questionnaires scores to ensure that affective ratings themselves were not linked to symptom severity, especially given that online populations are known to have increased mental illness severities (Chandler & Shapiro, 2016; Shapiro et al., 2013). We measured anxiety and obsessive-compulsion which are core components of two mental health disorders that often utilize aversive learning paradigms in research (Duits et al., 2015; D. J. Hauser & Schwarz, 2016). A prior in-lab rating study only collected state anxiety symptom data (Yang et al., 2018), which was akin to the severities of our current online sample. Overall, we found no significant relationship between the affective ratings and the psychiatric scores of the participant pool. Importantly, we also observed that the reliability of the ratings was not affected by the psychiatric symptoms, further supporting the use of these sounds in paradigms to study these illnesses. We acknowledge that as our results were drawn from a general population sample, its applicability to diagnosed patients may differ, but there is growing evidence that mechanisms captured with psychopathological variation in online general population findings are clinically relevant (Gillan et al., 2016, 2019).

There were some limitations to our study. Though we attained a certain degree of experimental control with our procedure, the ability to monitor sound presentation quality delivered to participants is still limited. Commercially available headphones may vary in their frequency response, and we were not able to check if participants had adjusted their system sound settings or had taken their headphones off after the screening test. Moreover, owing to security features of web browsers, it is impossible to track any computer system sound setting information—thus we cannot record the objective perceived sound intensities experienced by our participants. Secondly, the headphone screening test we utilized (Woods et al., 2017) relies on the ability of headphones to present separate signals to the two ears. As such, loudspeaker systems that broadcast a single channel/output of the combined audio could contribute to false positives. For future studies, recent developments of alternative headphone tests may help to resolve this issue (Milne et al., 2020). Despite the limitations, our findings show that affective responses are robust and can be evoked online to a level comparable with those collected in more controlled, in-lab environments.

Given that web-based crowdsourcing will become increasingly common for research, it is only fruitful to enable more paradigms to be translated to online platforms. While the degree of sound presentation control available may not be ideal, many web-induced limitations in other domains such as imprecisions in measuring reaction times (Bridges et al., 2020; Plant, 2016) have turned out to not compromise data substantially (Crump et al., 2013; Klein et al., 2014). Our findings present evidence that sounds presented through a web-based platform can evoke reliable affective ratings, which support the translation of affective audio-based paradigms to be conducted online.

## Data availability

Dataset and analysis code to reproduce all figures are freely available online at https://github.com/seowxft/audio-pilot-analysis.

## Acknowledgements

The authors thank Sijia Zhao for the initial discussion of the study. We also thank Felix Bacigalupo and Jayne Morris for sharing sounds utilized in the audio array.

This research was funded by a Wellcome Sir Henry Dale Fellowship (211155/Z/18/Z), a grant from the Jacobs Foundation (2017-1261-04), the Medical Research Foundation, and Brain & Behaviour Research Foundation NARSAD Young Investigator grant (27023) to TUH. The Max Planck UCL Centre for Computational Psychiatry and Ageing Research is a joint initiative supported by UCL and the Max Planck Society. The Wellcome Centre for Human Neuroimaging is supported by core funding from the Wellcome Trust (203147/Z/16/Z).

For the purpose of Open Access, the authors have applied a CC-BY public copyright license to any author accepted manuscript version arising from this submission.

