## Supplementary file 1 for "Reliability of web-based affective auditory stimulus presentation"

*for*

**Supplemental Methods**

***Age and gender.*** We examined the influence of age (*Age*, z-scored) and gender (*Gender*, as factor) on the affective ratings (*ScaleRating*) and questionnaire scores (*QuestionnaireScore*) with a mixed effect model and a linear regression model respectively. As gender assigned as ‘Other’ was underpowered (n = 1), we excluded that participant in these analyses that included gender as a covariate. For psychiatric scores, males tended to have lower state anxiety (*β* = -1.96, *SE* = 0.43, *p* < 0.001) and obsessive-compulsive symptom severity (*β* = -1.10, *SE* = 0.44, *p* = 0.01) than females, but no difference in trait anxiety scores (*β* = 0.75, *SE* = 0.45, *p* = 0.09). All psychiatric symptom scores overall decreased with age (STAI-Y1: *β* = -0.18, *SE* = 0.04, *p* < 0.001, STAI-Y2: *β* = -0.21, *SE* = 0.03, *p* < 0.001, OCI-R: *β* = -0.30, *SE* = 0.04, *p* < 0.001). As for the affective ratings, we found no age or gender difference (all *p*s > 0.20) nor interaction effects with of age/gender with volume on affective ratings (all *p*s > 0.12).

**Supplemental Figures**

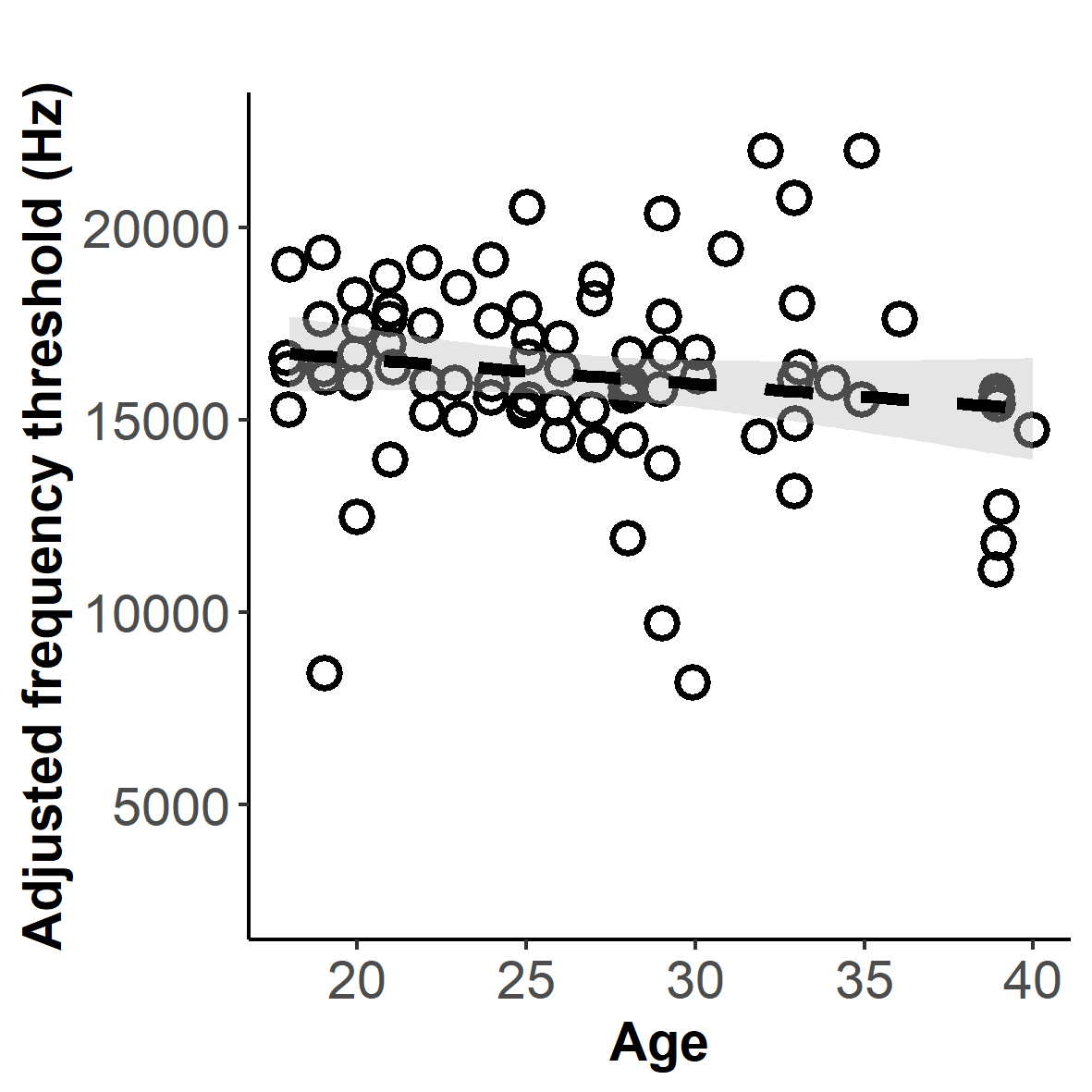

**Supplementary Fig. S1.** Relationship between final adjusted frequency threshold level (see *Sound frequency calibration*) and age. Sensitivity to high frequencies range tends to decline with age (Moore et al., 2014), thus we restricted our participant age range to 18 to 40 years old. There was no evidence of attenuated frequency sensitivity ability as age increased in our sample (*r* = -0.15, *p* = 0.16). Circles indicate the chosen frequency, dotted line indicates linear relationship and grey area indicates 95% confidence interval.

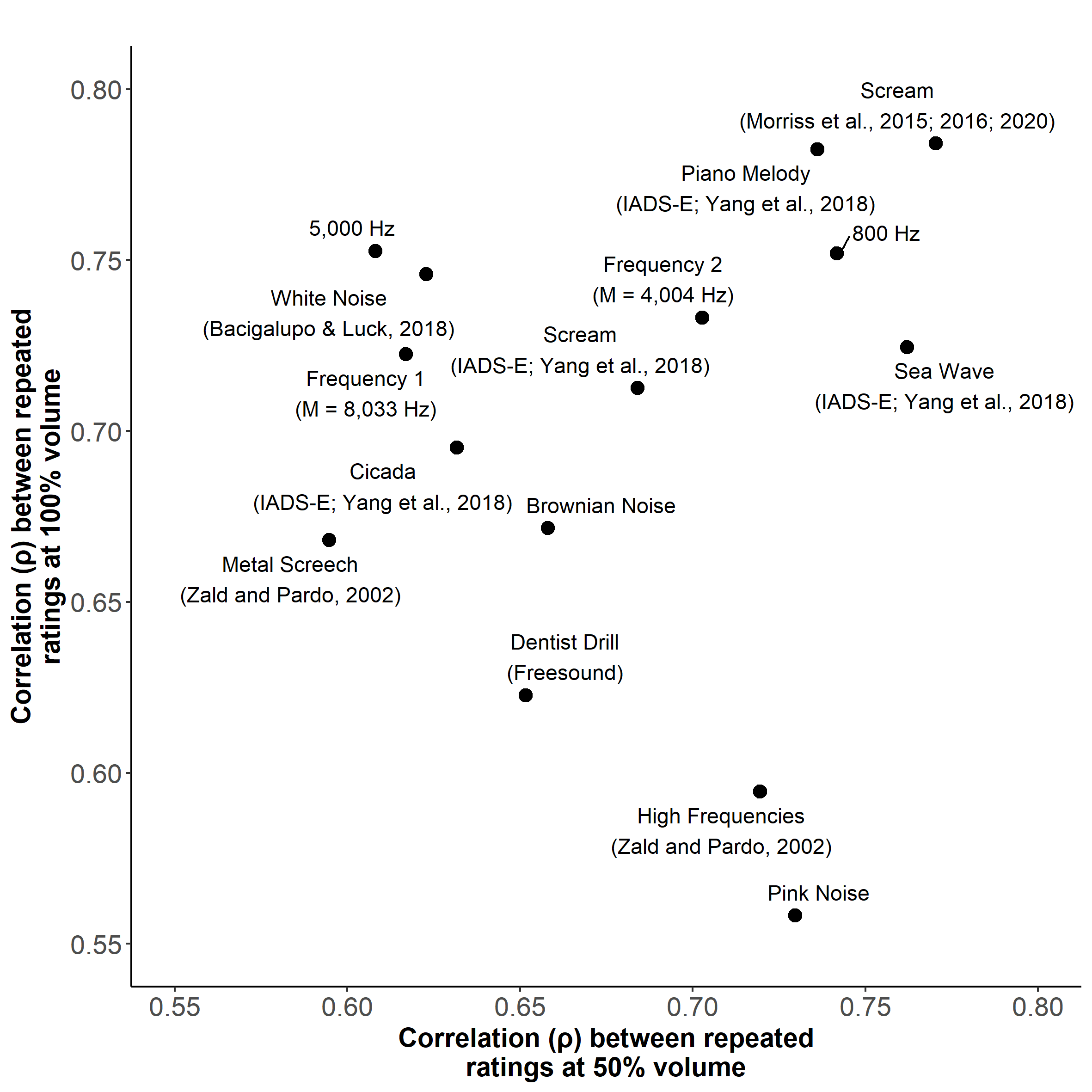

**Supplementary Fig. S2.** Sound-specific test-retest reliability. For each sound, ratings between repeated sound presentations at their specified volume levels were compared with Spearman’s correlation estimate *ρ*. Mean of the reliability estimates across all sounds for both volume levels were *ρ* ≥ 0.68, indicating that all sounds were generally able to reliably induce affective ratings.

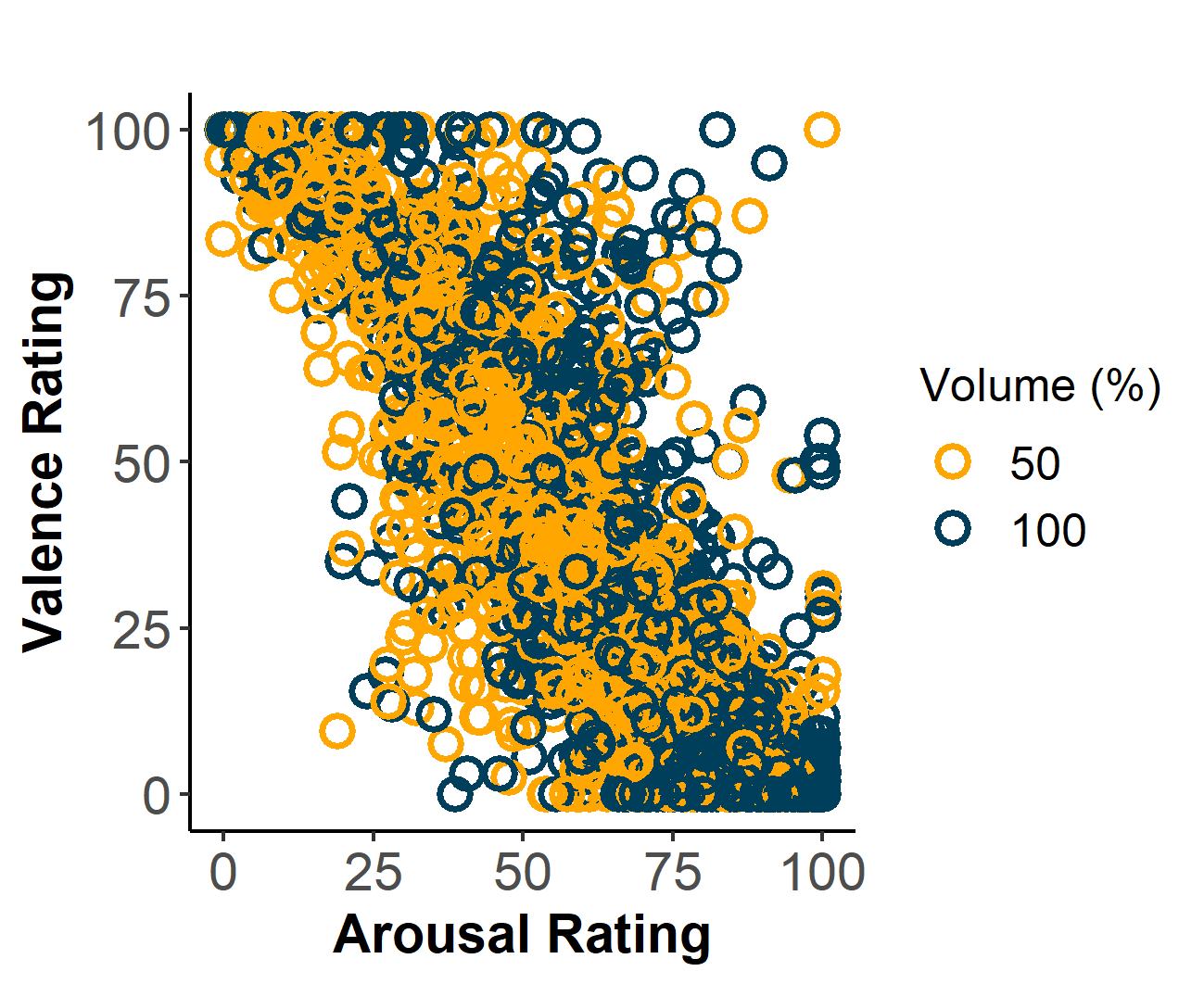

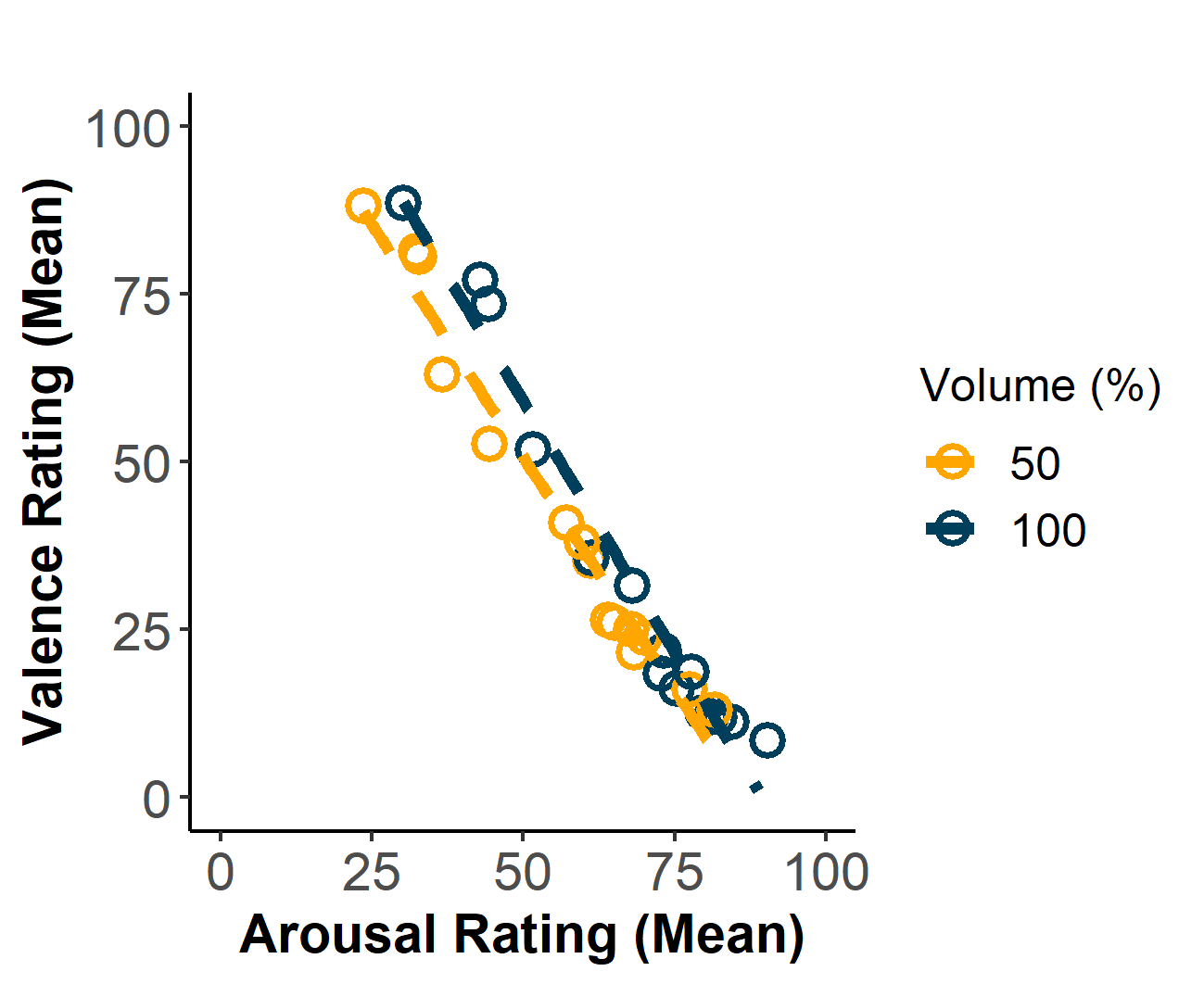

**Supplementary Fig. S3.** Scatterplot of valence versus arousal ratings. Left inset shows the ratings for all sound presentations while the right inset shows the mean rating across all participants for each sound. High arousal ratings (awakeness) were generally accompanied by low valence ratings (unpleasant), indicating a negative relationship. Circles represent the affective rating/mean affective rating in their respective graph. Dashed lines in the right graph depict the relationship between ratings in the two affective dimensions (Spearman: *ρ*: = -0.83, *p* < 0.001).

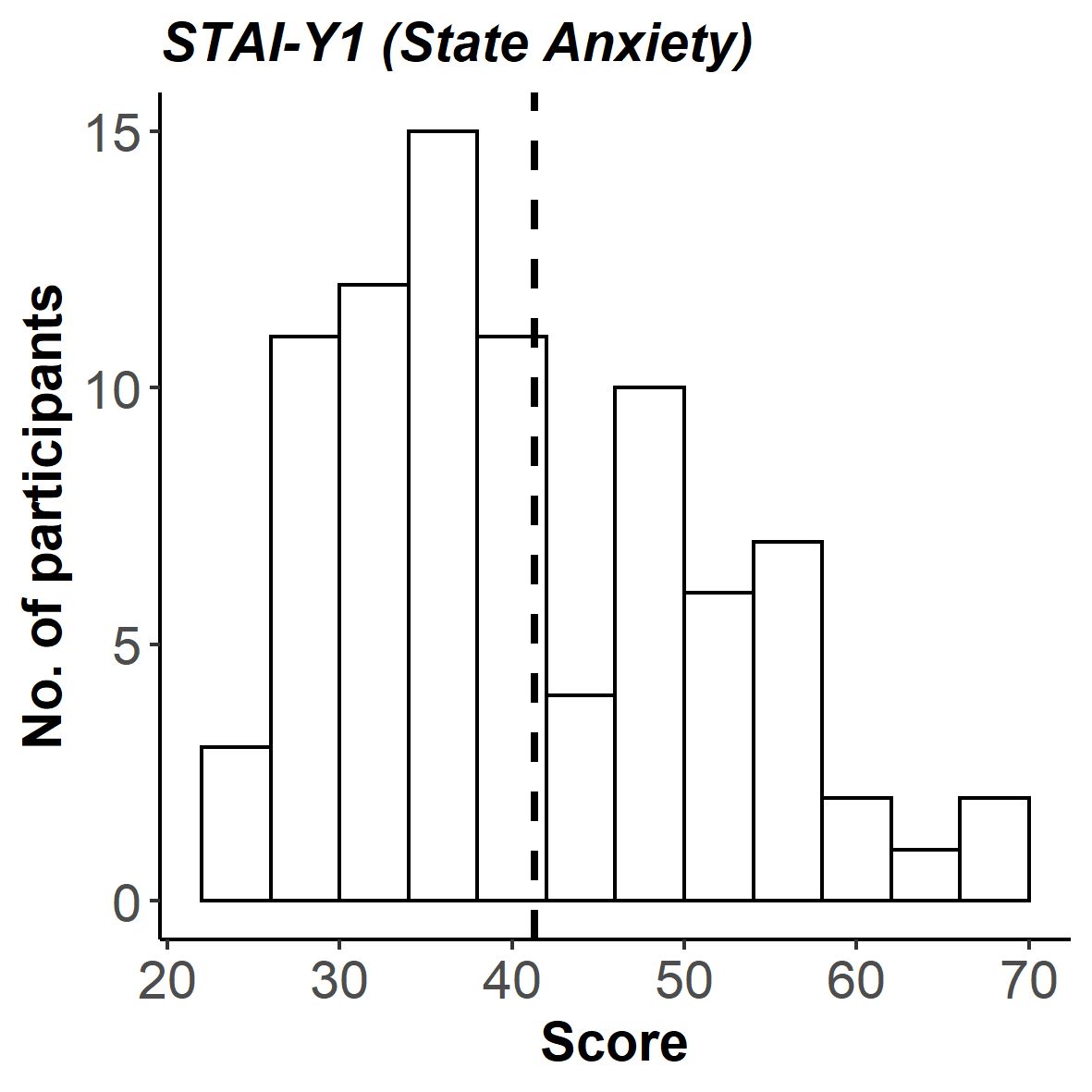

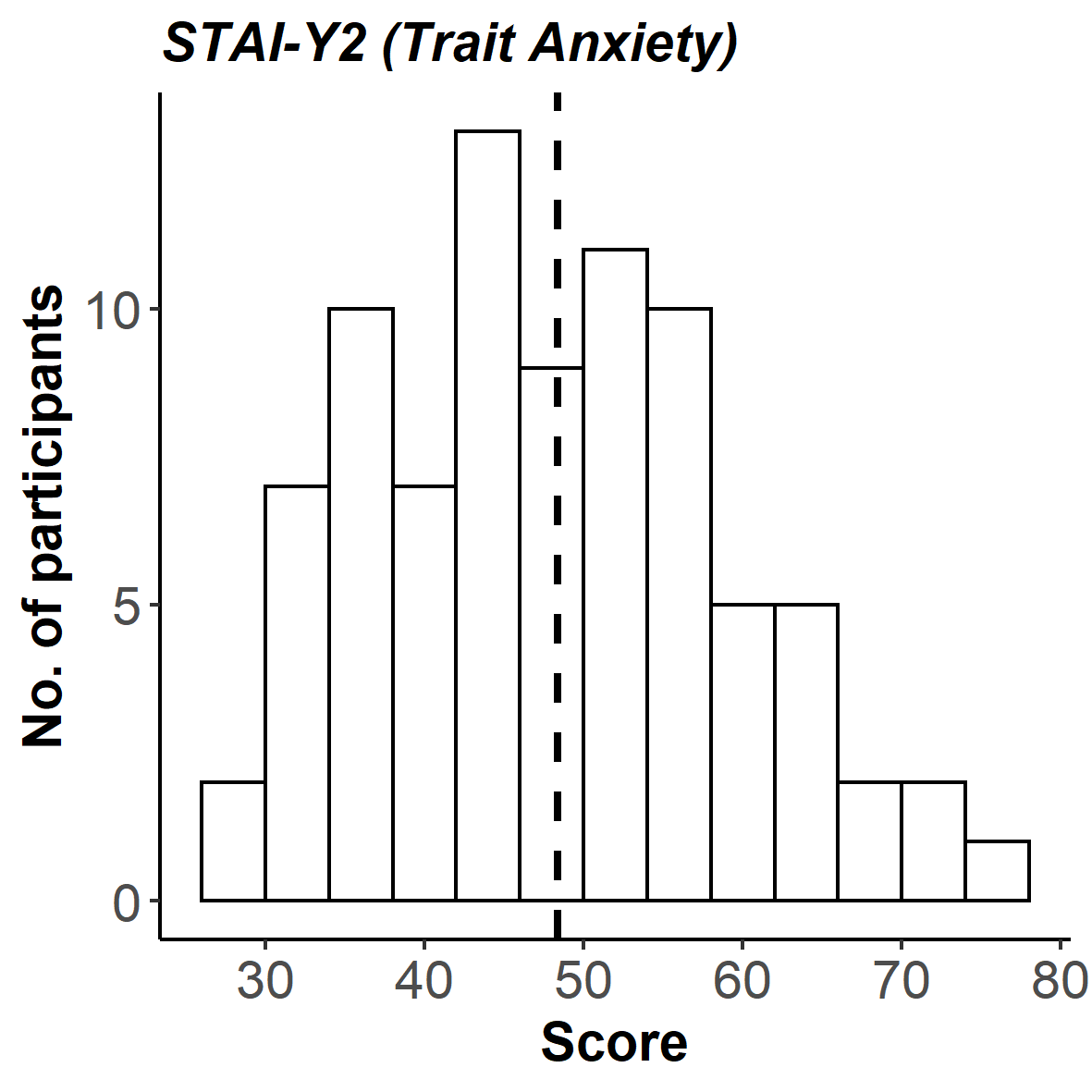

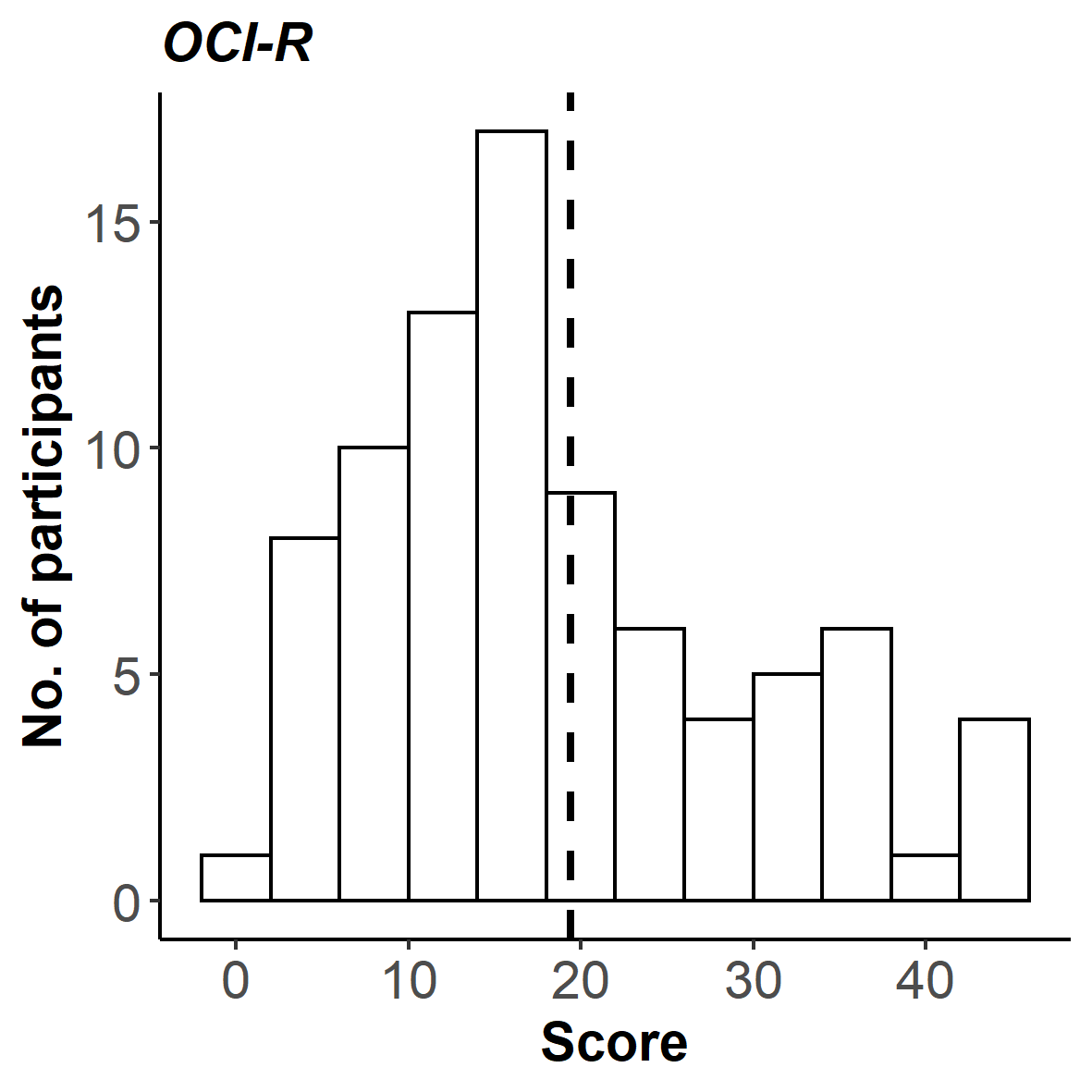

**Supplementary Fig. S4.** Histograms of questionnaire score distributions for state (STAI-Y1) and trait (STAI-Y2) anxiety, as well as obsessive-compulsive symptoms (OCI-R). Dashed lines indicate mean total score across all participants for the respective questionnaire: for the STAI-Y1: *M* = 41.32 (*SD* = 10.80), for the STAI-Y2: *M* = 48.39 (*SD* = 11.13) and for the OCI-R: *M* = 19.38 (*SD* = 11.36).

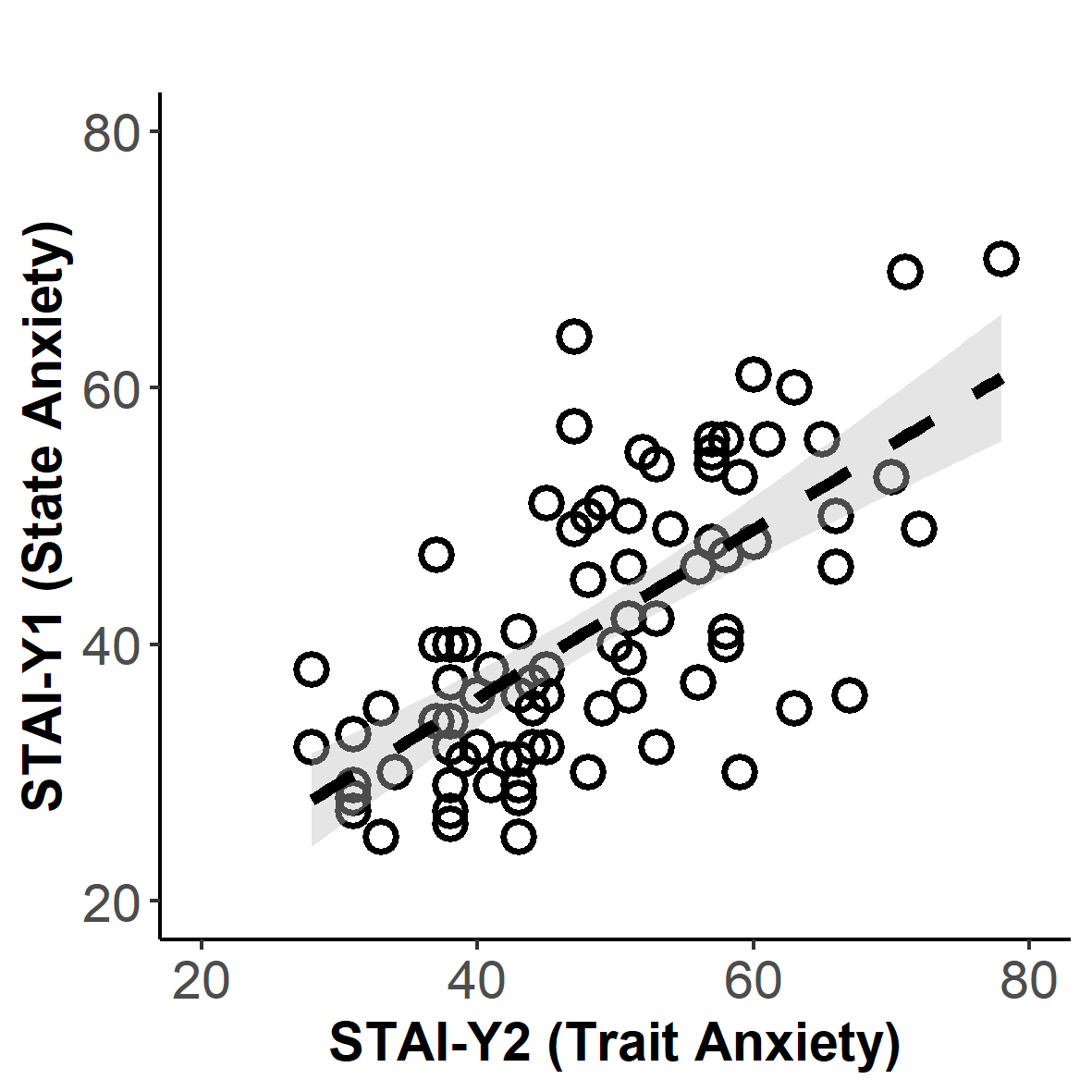

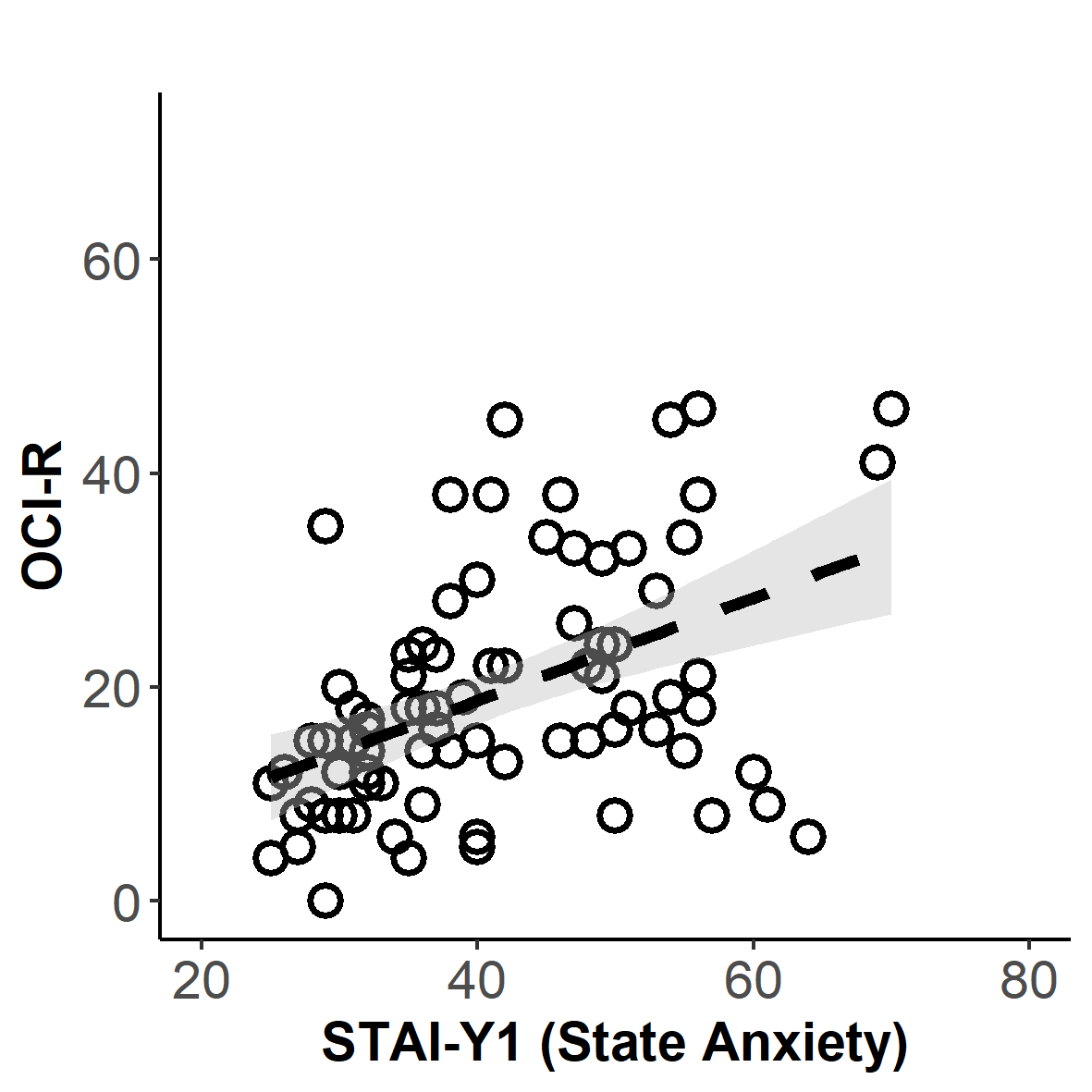

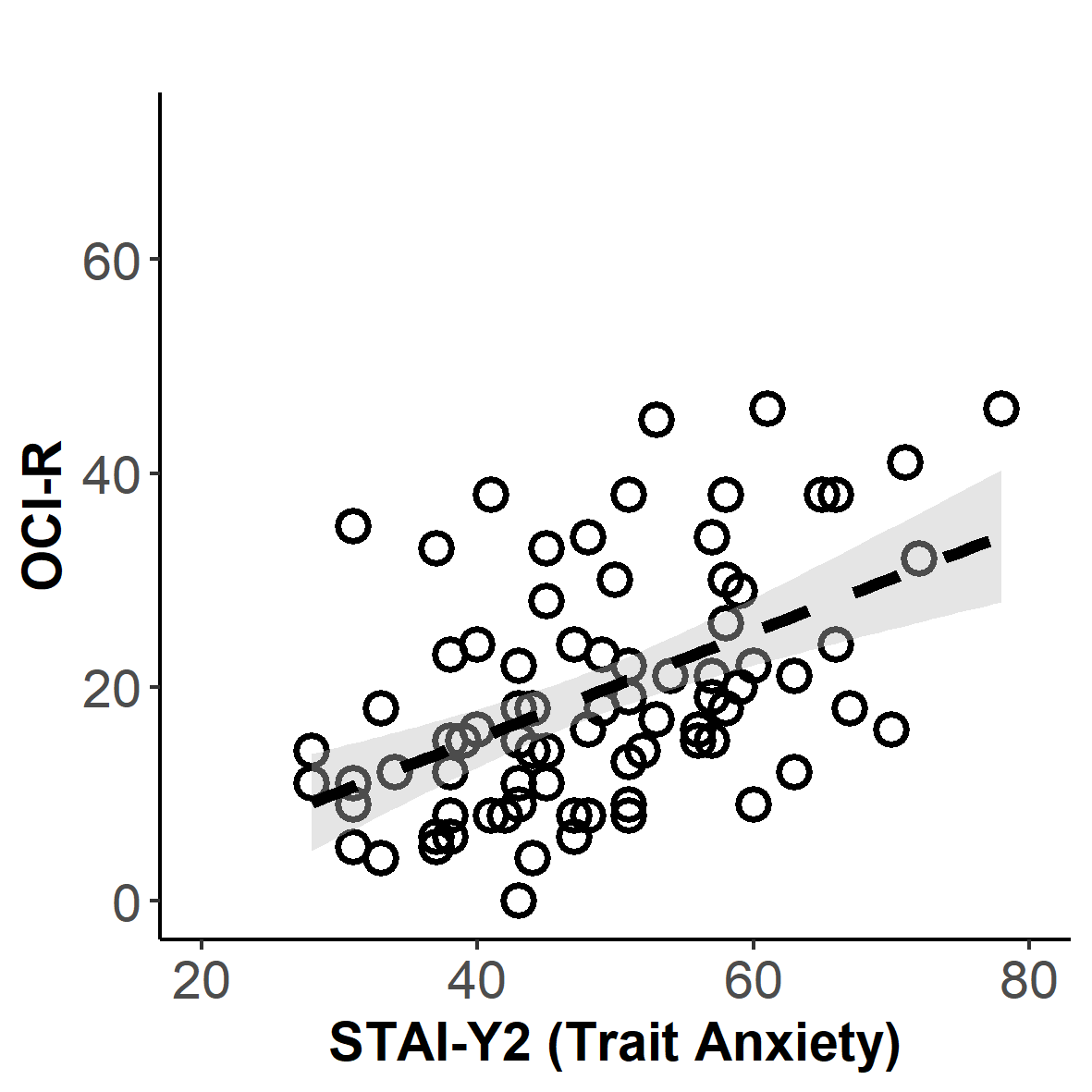

**Supplementary Fig. S5.** Relationship between psychiatric symptom scores. All questionnaires scores were positively correlated, between STAI-Y1 and STAI-Y2 (*r* = 0.68, *p* < 0.001), STAI-Y1 and OCI-R (*r* = 0.46, *p* < 0.001) and OCI-R and STAI-Y2 (*r* = 0.49, *p* < 0.001). Circles represent questionnaire total score per participant, dashed lines indicate linear relationship and grey areas indicate 95% confidence interval.

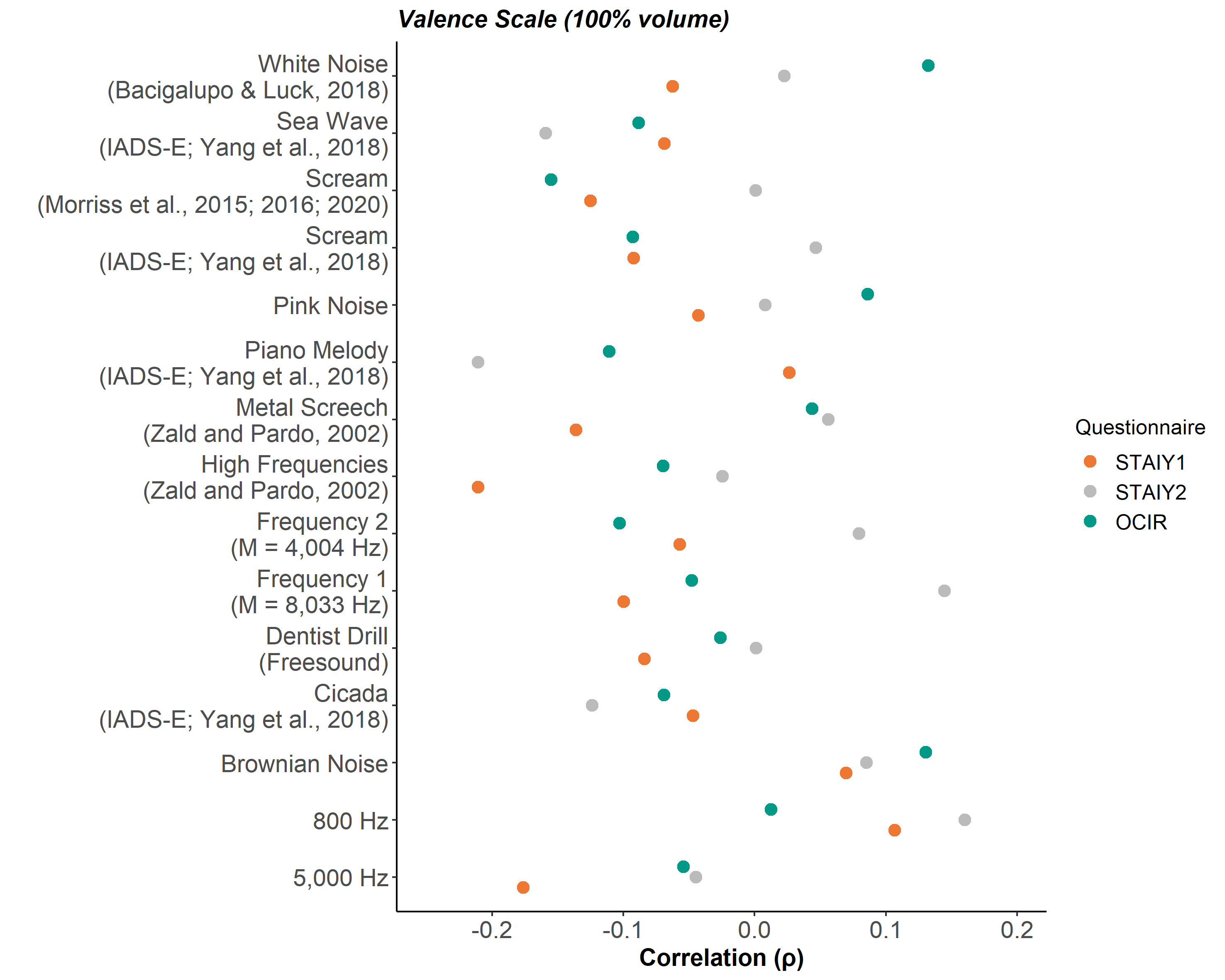

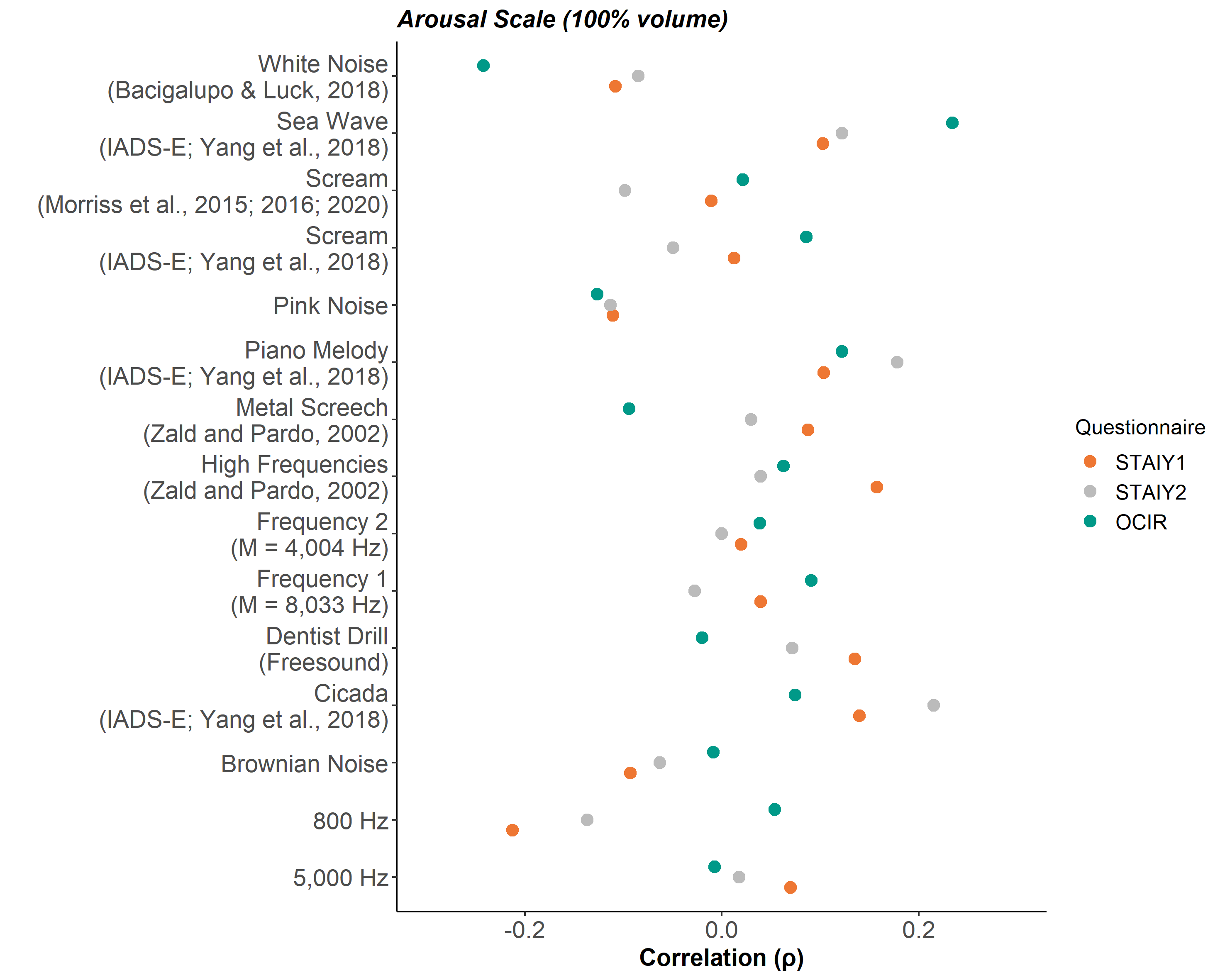

**Supplementary Fig. S6.** Sound-specific psychiatric symptom score correlation. For each individual sound played at 100% volume, ratings for all participants were correlated with each of the individuals’ total scores for STAI-Y1, STAI-Y2 and OCI-R separately. Majority of the relationships were non-significant, except for a few sounds in their arousal ratings where (i) individuals who scored high on the OCI-R tended to rate White Noise as less arousing (*ρ =* -0.24, *p* = 0.03, uncorrected) and Sea Wave as more arousing (*ρ =* 0.23, *p* = 0.03, uncorrected) and (ii) individuals who scored high on the STAI-Y2 tended to rate Cicada as more arousing (*ρ =* 0.23, *p* = 0.03, uncorrected).
